# Weighted likelihood inference of genomic autozygosity patterns in dense genotype data

**DOI:** 10.1101/177352

**Authors:** Alexandra Blant, Michelle Kwong, Zachary A. Szpiech, Trevor J. Pemberton

## Abstract

**Background:** Genomic regions of autozygosity (ROA) arise when an individual is homozygous for haplotypes inherited identical-by-descent from ancestors shared by both parents. Over the past decade, they have gained importance for understanding evolutionary history and the genetic basis of complex diseases and traits. However, methods to detect ROA in dense genotype data have not evolved in step with advances in genome technology that now enable us to rapidly create large high-resolution genotype datasets, limiting our ability to investigate their constituent ROA patterns.

**Results:** We report a weighted likelihood approach for identifying ROA in dense genotype data that accounts for autocorrelation among genotyped positions and the possibilities of unobserved mutation and recombination events, and variability in the confidence of individual genotype calls in whole genome sequence (WGS) data. Forward-time genetic simulations under two demographic scenarios that reflect situations where inbreeding and its effect on fitness are of interest suggest this approach is better powered than existing state-of-the-art methods to detect ROA at marker densities consistent with WGS and popular microarray genotyping platforms used in human and non-human studies. Moreover, we present evidence that suggests this approach is able to distinguish ROA arising via consanguinity from ROA arising via endogamy. Using subsets of The 1000 Genomes Project Phase 3 data we show that, relative to WGS, intermediate and long ROA are captured robustly with popular microarray platforms, while detection of short ROA is more variable and improves with marker density. Worldwide ROA patterns inferred from WGS data are found to accord well with those previously reported on the basis of microarray genotype data. Finally, we highlight the potential of this approach to detect genomic regions enriched for autozygosity signals in one group relative to another based upon comparisons of per-individual autozygosity likelihoods instead of inferred ROA frequencies.

**Conclusions:** This weighted likelihood ROA detection approach can assist population- and disease-geneticists working with a wide variety of data types and species to explore ROA patterns and to identify genomic regions with differential ROA signals among groups, thereby advancing our understanding of evolutionary history and the role of recessive variation in phenotypic variation and disease.

## Background

Genomic regions of autozygosity (ROA) reflect homozygosity for haplotypes inherited identical-by-descent (IBD) from an ancestor shared by both maternal and paternal lines. Common ROA are a source of genetic variation among individuals that can provide invaluable insight into how population history, such as bottlenecks and isolation, and "sociogenetic" factors, such as frequency of consanguineous marriage, influence genomic variation patterns. Population-genetic studies in worldwide human populations over the past decade have found ROA ranging in size from tens of kb to multiple Mb to be ubiquitous and frequent even in ostensibly outbred populations [1–28] and to have a non-uniform distribution across the genome [7,10,13,18] that is correlated with spatially variable genomic properties [2–4,18] creating autozygosity hotspots and coldspots [18]. ROA of different sizes have different continental patterns both with regards to their total lengths in individual genomes [12,18,22,24,26–28] and in their distribution across the genome [18] reflecting the distinct forces generating ROA of different lengths. Studies of ROA in the genomes of ancient hominins [29–31] and early Europeans [32] have provided unique insights into the mating patterns and effective population sizes of our early forbearers. In non-humans, ROA patterns have provided insights into the differential histories of woolly mammoth [33], great ape [34,35], cat [36], canid [37–42], and bird [43] populations, while in livestock breeds they have provided insights into their origins, relationships, and recent management [42,44–61] and the lasting effects of artificial section [58,61–73], as well as informed the design of ongoing breeding [74,75] and conservation [47,57,76] programs [77].

In contemporary human populations, increased risks for both monogenic [78–82] and complex [83–90] disorders as well as increased susceptibility to some infectious diseases [91–93] have been observed among individuals with higher levels of parental relatedness. While the association between parental relatedness and monogenic disease risk has been known for more than a century [94], observations with complex and infectious diseases potentially reflect elevated levels of autozygosity as a consequence of prescribed and unintentional inbreeding [95] that enrich individual genomes for deleterious variation carried in homozygous form [96,97]. Indeed, genomic autozygosity levels have been reported to influence a number of complex traits, including height and weight [98–101], cognitive ability [101–103], blood pressure [104–111], and cholesterol levels [111], as well as risk for complex diseases such as cancer [84,85,112–116], coronary heart disease [84,117–119], amyotrophic lateral sclerosis (ALS) [120], and mental disorders [121,122]. These observations are consistent with the view that variants with individually small effect sizes associated with complex traits and diseases are more likely to be rare than to be common [123–126], are more likely to be distributed abundantly rather than sparsely across the genome [9,127], and are more likely to be recessive than to be dominant [9,128]. Recent studies investigating ROA and human disease risk have identified both known and novel loci associated with standing height [129], rheumatoid arthritis [130], early-onset Parkinson’s disease [131], Alzheimer’s disease [132,133], ALS [120], schizophrenia [4,134], thyroid cancer [116], and Hodgkin lymphoma [115,135]. Thus, just as ROA sharing among affected individuals has facilitated our understanding of the genetic basis of monogenic disorders [136] in both inbred [137–140] and more outbred [141–143] families, it also represents a potentially powerful approach with which to further our understanding of the genetic etiology of complex disorders [144] of major public health concern worldwide.

In both population- and disease-genetic studies, ROA are frequently inferred from runs of homozygous genotypes (ROH) present in genome-wide single nucleotide polymorphism (SNP) data obtained using high-density microarray platforms [145]. A popular program for ROH identification is *PLINK* [146], which uses a sliding window framework to find stretches of contiguous homozygous genotypes spanning more than a certain number of SNPs and/or kb, allowing for a certain number of missing and/or heterozygous genotypes per window to account for possible genotyping errors. While a number of more advanced ROA identification approaches have been proposed [147,148], a recent comparison found the *PLINK* method to outperform these alternatives [149]. We recently proposed to detect ROA using a sliding-window framework and a logarithm-of-the-odds (*LOD*) score measure of autozygosity [1,150] that offers several key advantages over the *PLINK* method [18]. First, it is not reliant on fixed parameters for the number of heterozygous and missing genotypes when determining the autozygosity status of a window, instead incorporating an assumed genotyping error rate, making it more robust to missing data and genotyping errors. Second, it incorporates allele frequencies in the general population to provide a measure of the probability that a given window is homozygous by chance, allowing homozygous windows to be distinguished from autozygous windows. These important advances would be expected to provide greater sensitivity and specificity for the detection of ROA in high-density SNP genotype data, particularly in the presence of the higher and more variable genotype error rates in next-generation sequence (NGS) data [151,152].

A shortcoming of the *LOD* method is that correlations between SNPs within a window that occur as a consequence of linkage disequilibrium (LD) are ignored, leading to overestimation of the amount of information that is available in the data and potentially false-positive detection of autozygosity signals. In addition, the *LOD* method does not account for the possibility of recent recombination events onto very similar haplotype backgrounds that might give the appearance of autozygosity when paired with a non-recombined haplotype [153]. Such a scenario would, for example, arise when ROA are detected in microarray-based genotype data that comprises information at only a limited set of positions within a genomic interval and is therefore blind to unobserved genetic differences that make the apparently identical haplotypes distinct.

Here, we report an improved *LOD*-based ROA detection method that accounts for the non-independence between SNPs and the likelihoods of unobserved mutation and recombination events within a window. We compare the performance of this new method against the original *LOD* method as well as a newly reported method implemented in the *BCFtools* software package [154] in simulated genetic datasets. We then evaluate how ROA inference is influenced by the source and density of interrogated markers using the 26 worldwide human populations included in Phase 3 of The 1000 Genomes Project [155], considering the entire whole-genome sequence (WGS) dataset as well as subsets representing SNPs present in the exome and included on two commonly used Illumina BeadChips. We show population differences in genome-wide ROA patterns inferred from WGS data using our improved *LOD*-based method recapitulate those observed in our earlier BeadChip-based study that used the original *LOD* method [18]. Finally, we highlight the unique ability of our improved *LOD*-based method to identify genomic regions enriched for autozygosity signals in one group relative to another without first inferring ROA through the direct comparison of weighted *LOD* scores, finding nine regions that significantly differ in the strength of their autozygosity signals between apparent subgroups within the Asian Indian Gujarati, Punjabi, and Telugu populations. Our improved ROA detection method will assist population- and disease-geneticists working with a wide variety of data types and species to explore ROA patterns and to identify genomic regions with differential ROA signals, thereby facilitating our understanding of the role of recessive variants in phenotypic variation and disease.

## Results

### Weighted likelihood autozygosity estimator

We previously reported an ROA detection approach that was based on a number of earlier methods [1,150] in which a likelihood-based autozygosity estimator is applied in a sliding window framework where window size is defined as a fixed number of SNPs [18]. In this approach, within window *w* in individual *i* from population *j*, the *LOD* score of autozygosity is calculated across the *K* SNP markers within window *w*, where we observe genotype *G*_*k*_ at the *k*^*th*^ SNP that has state *X*_*k*_, which equals 1 if the SNP is autozygous and 0 otherwise.

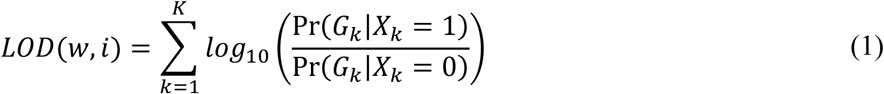

The per-SNP likelihoods of autozygosity and non-autozygosity are based on Hardy-Weinberg proportions (**Table 1**) and include population-specific allele frequencies and an assumed rate of genotyping errors and mutations ε. Missing genotypes are ignored in this algorithm; that is, they have a log-likelihood of zero. The log-likelihood of autozygosity for homozygous SNPs is positive and decreases exponentially as a function of allele frequency (Additional File 1: **Figure S1A**). The log-likelihood of autozygosity for heterozygous SNPs is instead negative and equal to log_10_ (ε), thus acting as a penalty for the presence of heterozygous genotypes within a window.

**Table 1.**
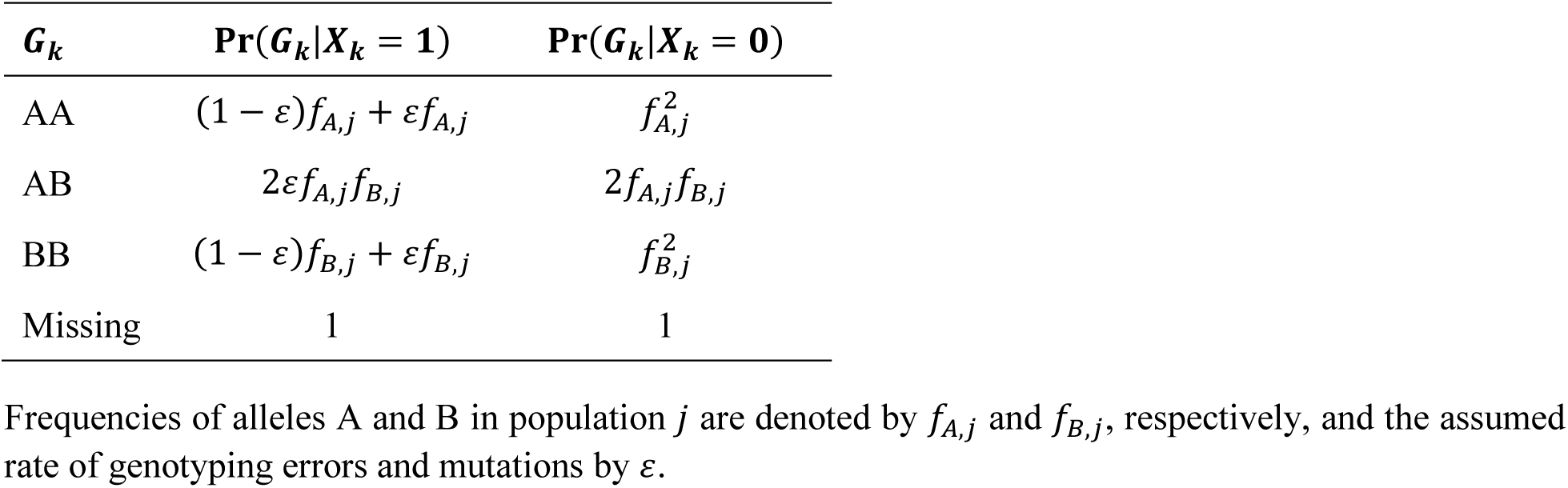
Per-SNP likelihoods of autozygosity and non-autozygosity.

To address the apparent shortcomings of the *LOD* score method, we developed a weighted *LOD*-based method (*wLOD*) that accounts for non-independence among SNPs and the probabilities of recombination and mutation within window *w*.

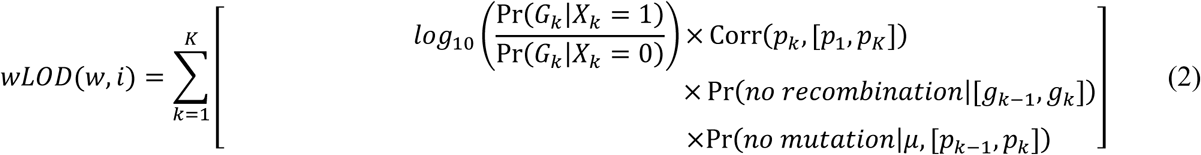

Here, we adapt the approach of Chen *et al*. [156] to incorporate LD information into the *wLOD*(*w*, *i*) estimator, weighting the log-likelihood of SNP *k* by the reciprocal of the sum of pairwise LD between SNP *k* and all other SNPs within window *w* calculated as

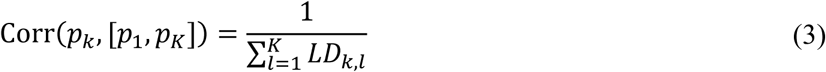

and bounded in the interval [1/*K*,1]. An intuitive explanation for this correction is that when a number of SNPs are highly correlated they provide redundant information. By weighting the log-likelihood for SNP *k* as a function of its correlation with all other SNPs within window *w* it contributes only the unique autozygosity information it possesses to w*LOD*(*w*, *i*).

LD reflects historical recombination and mating patterns in a population and is largely insensitive to the effects of mating patterns within the last few generations that can, through recombination events onto very similar haplotype backgrounds, lead to false-positive autozygosity signals [153]. Thus, we also weight the log-likelihood of SNP *k* by the probability of no recombination events having occurred within the genomic interval bounded by SNP *k* − 1 and SNP *k* in the last *M* generations, calculated based upon their genetic map position *g* (in Morgans) as previously described [10,157]

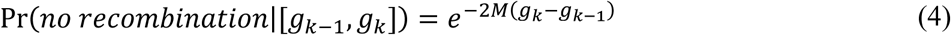

In a population-genetic context, *M* can be set based upon effective population size estimates and the probability that a pair of individuals share a common ancestor *M* generations in the past [158], while in a disease-genetic context *M* can instead be set based on known relationships between affected individuals.

Finally, we account for the potential presence of unobserved genetic differences within the genomic interval bounded by SNP *k* − 1 and SNP *k* by weighting the log-likelihood of SNP *k* by the probability of no unobserved mutation events having occurred within the genomic interval in the last *M* generations, calculated based upon their physical map position *p* (in bp) and a per-base mutation rate μ using an approach adapted from MacLeod *et al*. [159]

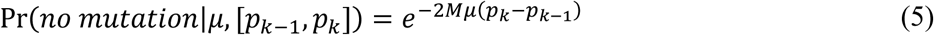

As evident in **Figure S1B** (Additional File 1), the recombination and mutation weightings reduce the log-likelihood of SNP *k* as a function of its distance from SNP *k* − 1. It can also be seen that as *M* decreases the magnitude of the change in the weighting with increasing distance also decreases; thus, *wLOD* scores in populations with small effective population sizes or in disease studies where affected individuals share a more recent common ancestor (smaller *M*) will be adjusted to a lesser extent than those with larger effective population sizes or where affected individuals share a more distant common ancestor (larger *M*).

### Properties of the *wLOD* estimator

We investigated the properties of the *wLOD* estimator using The 1000 Genomes Project Phase 3 dataset that provides phased genotypes for 84,801,880 genetic variants discovered using a low-coverage WGS approach in 2,436 unrelated individuals from 26 worldwide human populations (**Table 2**) [155]. To approximate a typical microarray-based SNP genotyping study, we first developed a subset of this dataset that contained 2,166,414 autosomal SNPs that are present on the popular Illumina HumanOmni2.5-8 BeadChip (“Omni2.5 dataset” henceforth). In all analyses, μ was set to 1.18×10^−8^ [160] and ε was set to 4.71×10^−4^, the average rate of discordance across samples between genotypes in our Omni2.5 dataset and those obtained for 1,693 of the 2,436 individuals directly with the Illumina HumanOmni2.5 BeadChip [155]. Unless otherwise stated, *M* was set to seven, a conservative value broadly reflecting the average of effective population size estimates for populations included in The 1000 Genome Project [155,158,161]. Window size was varied in an arbitrary interval [*K*_0_, *K*_*n*_] in which *K* is increased in 10 SNP increments (i.e. *K*_*n*_ = *K*_0_ + [10×*n*]).

**Table 2.** Populations included in Phase 3 of The 1000 Genomes Project.

| Population |  | Geographic region | *N* | Consanguinity<sup>a</sup> |  |
| --- | --- | --- | --- | --- | --- |
| ID | Name |  |  | Frequency | Reference(s) |
| ESN | Esan | Africa | 94 | – | – |
| GWD | Gambian | Africa | 109 | – | – |
| LWK | Luhya | Africa | 96 | – | – |
| MSL | Mende | Africa | 80 | – | – |
| YRI | Yoruban | Africa | 107 | 51.20% | [250] |
| GBR | British | Europe | 89 | 0.40% | [251] |
| CEU | European American | Europe | 97 | 0.20% | [252] |
| FIN | Finnish | Europe | 98 | 0.17% | [253] |
| IBS | Iberian | Europe | 107 | 1.99% | [254-258] |
| TSI | Toscani | Europe | 104 | – | – |
| BEB | Bengali | Central/South Asia | 84 | 5.00% | [173] |
| GIH | Gujarati | Central/South Asia | 101 | 4.90% | [173] |
| PJL | Punjabi | Central/South Asia | 96 | 0.90% | [173] |
| STU | Sri Lankan Tamil | Central/South Asia | 96 | 38.20% | [173] |
| ITU | Telugu | Central/South Asia | 101 | 30.80% | [173] |
| CDX | Dai | East Asia | 92 | 21.30% | [172] |
| JPT | Japanese | East Asia | 103 | 4.80% |  |
| KHV | Kinh | East Asia | 94 | – | – |
| CHB | Northern Han | East Asia | 101 | 1.16% | [172,259,260] |
| CHS | Southern Han | East Asia | 102 | 3.43% | [172,260] |
| ASW | African American | Admixed | 55 | – | – |
| ACB | Afro-Caribbean | Admixed | 94 | – | – |
| CLM | Colombian | Admixed | 89 | 2.83% | [252,254,261] |
| MXL | Mexican American | Admixed | 62 | 0.80% | [252,254] |
| PEL | Peruvian | Admixed | 84 | 1.90% | [252,261,262] |
| PUR | Puerto Rican | Admixed | 101 | 3.30% | [257] |

<sup>a</sup>Consanguinity frequencies were obtained from http://www.consang.net.

The genome-wide distribution of *wLOD* scores for all windows in the Omni2.5 dataset is bimodal and centered around 0 (**Figure 1A**), with *wLOD* scores under the left-hand mode favoring the hypothesis of non-autozygosity, whereas those under the right-hand mode favor the autozygosity hypothesis. The area under the right-hand mode decreases as a function of window size as ROA are progressively covered by fewer but longer windows. In addition, while the location of the right-hand mode does not change appreciably with window size, there is a noticeable shift toward lower *wLOD* scores in the left-hand mode with increasing window size, likely reflecting the larger number of heterozygous SNPs in non-autozygous compared with autozygous regions and their greater cumulative effect on *wLOD* scores with increasing window size. This shift progressively increases the distance between the non-autozygous and autozygous modes until either the autozygous mode disappears (**Figure 1B**) or the intermodal distance begins to decrease instead (Additional File 1: **Figure S2**), both potentially reflecting the point above which window lengths exceed those of the majority of ROA in the sample set. In this scenario, as window size increases autozygous windows increasingly overlap non-autozygous regions flanking shorter ROA leading them to encompass greater numbers of heterozygotes within these non-autozygous regions, deflating their *wLOD* scores. Whether the autozygous mode disappears or shifts toward lower *wLOD* scores is likely determined by the abundance of ROA and their levels of support in the sample set: sets with fewer ROA and ROA with generally lower *wLOD* scores lose their autozygous mode while those with large numbers and higher *wLOD* scores have it shift toward the non-autozygous mode. Nevertheless, the location of the minimum between the two modes does shift subtly toward higher *wLOD* scores with increasing window size, reflecting the expected increase in scores for autozygous windows as a function of the number of SNPs within the window. The periodicity observed in the genome-wide score distribution of the original *LOD* estimator [18] is absent with the *wLOD* estimator, indicating that this property was a reflection of LD among SNPs within the window.

**Figure 1.**
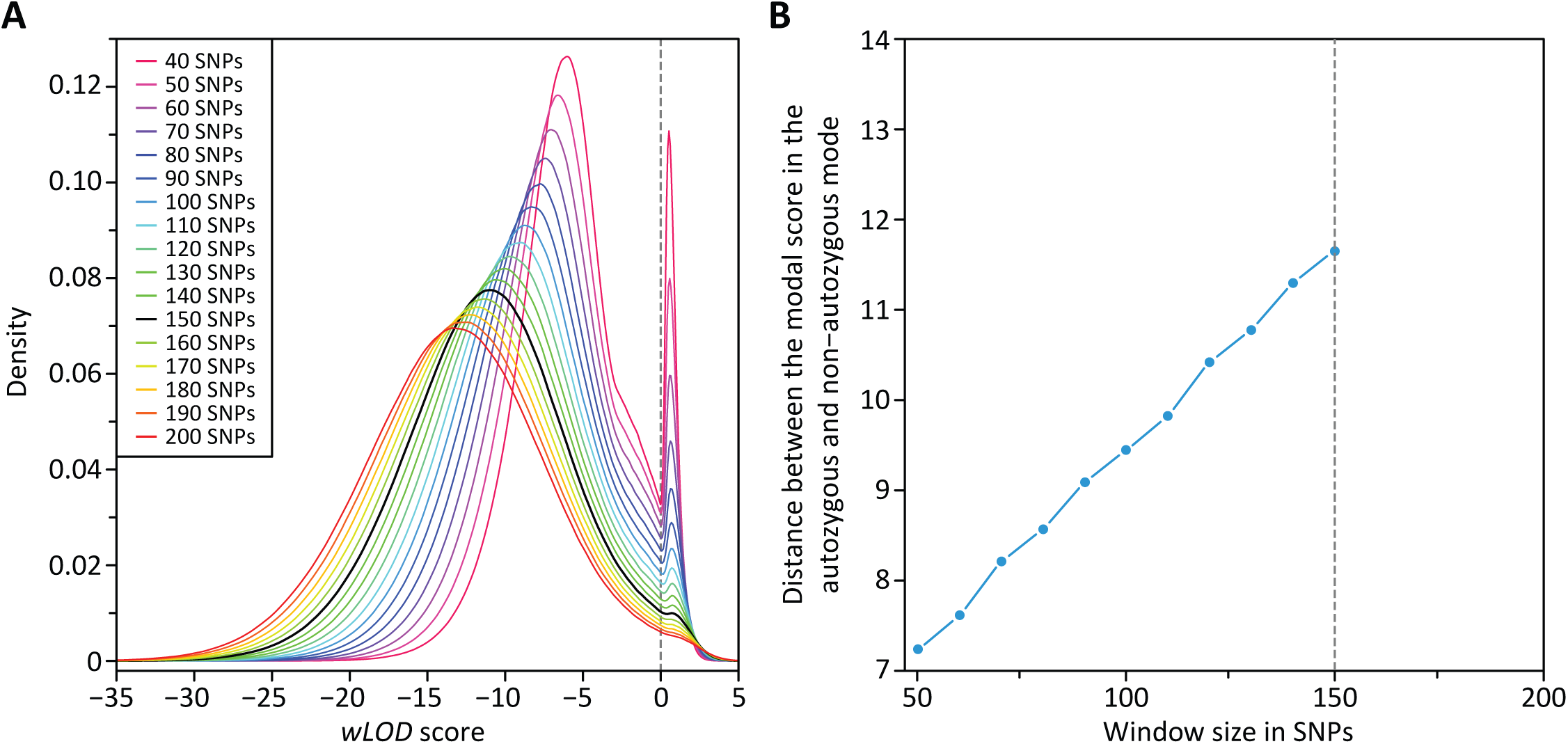
Distribution of genome-wide scores in European Americans. (**A**) Each line represents the Gaussian kernel density estimates of the pooled scores from all 97 individuals in the European American (CEU) population at window sizes between 40 and 200 SNPs in 10 SNP increments in the Omni2.5 dataset. The largest window size that produced a clear bimodal distribution (150 SNPs) is shown in black. (**B**) The change in intermodal distance with increasing window size in the CEU population. These patterns are representative of those observed in all other populations in the dataset.

To evaluate how the improvements incorporated into the *wLOD* estimator (equation 3) influence per-window scores as compared to the original *LOD* estimator (equation 1), we compared *wLOD* and *LOD* scores in the Omni2.5 dataset with a window size of 150 SNPs (**Figure 2A**), the largest value that produced a clear bimodal *wLOD* score distribution in all populations. Across populations, per-window *wLOD* scores differed from their corresponding *LOD* scores by between −103.87 and 454.07 (**Figure 2B**) with the range and average of *wLOD* and *LOD* score differences increasing as a function of a population’s geographic distance from East Africa (*ρ*=0.8460 with *P*=5.029×10^−6^ and *ρ*=0.8846 with *P*=4.961×10^−7^, respectively), reflecting increasing LD [162,163] and decreasing genetic diversity [95,164–167]-leading to larger inter-SNP distances-with distance from Africa. Among the six admixed populations included in Phase 3 of The 1000 Genomes Project, those of mixed African and European ancestry (ACB and ASW) had smaller ranges and averages of *wLOD* and *LOD* score differences than those of mixed of Amerindian and European ancestry (CML, MXL, PUR, and PEL), consistent with the lower LD [168–170] and higher genetic diversity [167,171] of admixed African-European populations compared with Amerindian-European populations.

**Figure 2.**
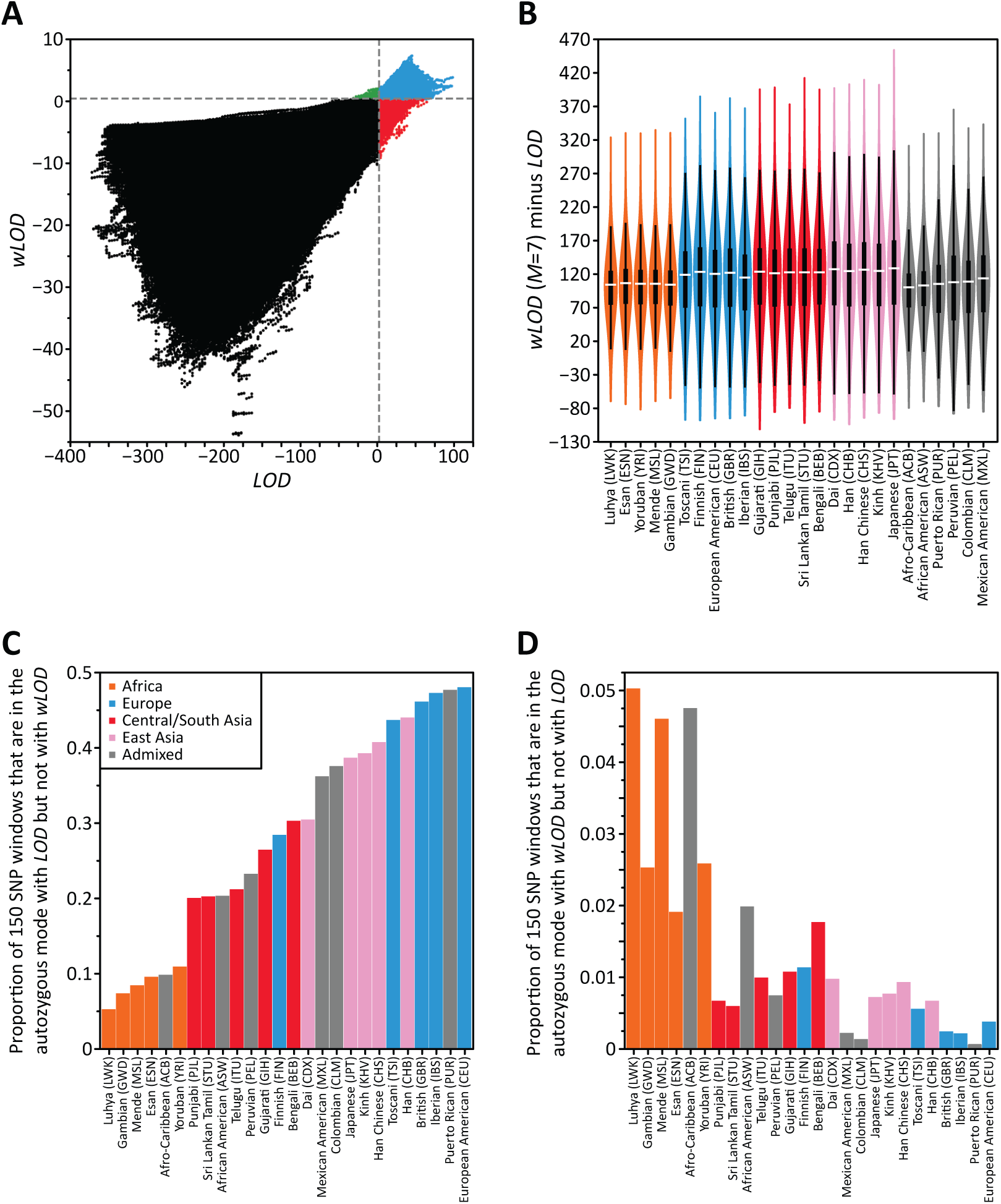
Difference in per-window scores between the and estimators. (**A**) Scatterplot comparing per-window and scores across all individuals in the European American (CEU) population at a window size of 150 SNPs in the Omni2.5 dataset. The 2,675,059 windows that were in the autozygous mode with both and are shown in blue. The 9,885 windows that were in the non-autozygous mode with but the autozygous mode with are shown in green. The 2,462,843 windows that were in the autozygous mode with but the non-autozygous mode with are shown in red. All windows that were in the non-autozygous mode with both and are shown in black. (**B**) Violin plots representing the change in per-window score between and across all individuals in each population for 150 SNP windows in the Omni2.5 dataset. Each ‘‘violin’’ contains a vertical black line (25%–75% range) and a horizontal white line (median), with the width depicting a 90°-rotated kernel density trace and its reflection, both colored by the geographic affiliation of the population [249]. Bar plots showing for each population the proportion of 150 SNP windows that are (**C**) in the autozygous mode with the estimator but are in the non-autozygous mode with the estimator or (**D**) in the non-autozygous mode with the estimator but are in the autozygous mode with the estimator.

Across populations, 5.15–47.93% of all windows in the right-hand “autozygous” mode with the *LOD* estimator were present in the left-hand “non-autozygous” mode with the *wLOD* estimator (**Figure 2C**) potentially reflecting false-positive autozygosity signals reported by the *LOD* estimator as a consequence of non-independence among homozygous SNPs that cumulatively give the mistaken impression of autozygosity. The proportion of windows was lowest in African populations and highest in most European populations, increasing incrementally through Central/South Asian and East Asian populations. This pattern can be explained by population differences in the location of the autozygous mode and its shift toward lower scores with the *wLOD* estimator. Modal *LOD* and *wLOD* scores in the autozygous mode are generally smallest and most similar in European populations and highest and least similar in African populations (Additional File 1: **Figure S3A**). Thus, for a given unit decrease in score between the *LOD* and *wLOD* estimators, an autozygous *LOD* window has a greater chance of transitioning to the non-autozygous *wLOD* mode in Europeans populations than in African populations. Consistent with this hypothesis, the magnitude of the difference between modal *LOD* and *wLOD* scores in the autozygous mode and the location of the minima between the autozygous and non-autozygous modes is significantly negatively correlated with the proportion of autozygous *LOD* windows that transition to the non-autozygous *wLOD* mode (*r*=-0.8654, *P*=1.156×10^−8^; Additional File 1: **Figure S3B**).

In contrast, across populations only 0.055–5.015% of all windows in the non-autozygous mode with the *LOD* estimator were present in the autozygous mode with the *wLOD* estimator (**Figure 2D**), potentially reflecting false-negative autozygosity signals reported by the *LOD* estimator as a consequence of heterozygotes in high LD with a larger number of homozygotes that, in one possibility, might reflect genotyping errors. The proportion of windows was highest in most African populations and lowest in most European populations, with broadly similar values observed in Central/South and East Asian populations. This pattern is the opposite of that observed with the putative false-positive windows above, and can also be explained by population differences in the location of the autozygous mode and its shift toward lower scores with the *wLOD* estimator. The addition of a single heterozygote to an autozygous window in the European populations has a greater chance of transitioning it from the autozygous to non-autozygous mode than in the African populations since the autozygous mode is situated much closer to the minima between the two modes (Additional File 1: **Figure S3**).

Overall, the much larger numbers of windows transitioning from the autozygous to the non-autozygous mode than vice versa between the *LOD* and *wLOD* estimators accords with the expectation that the *LOD* estimator frequently overestimates the amount of information available in the data leading it to falsely report autozygosity signals particularly in genomic regions with higher levels of LD, while it underestimates the amount of information much less frequently.

### Evidence of separate endogamic and consanguinity autozygosity signals in Asian Indians

In four of the five Asian Indian populations–Gujarati (GIH), Telugu (ITU), Punjabi (PJL), and Sri Lankan Tamil (STU)–as well as in the East Asian Dai (CDX) population, as window size increased a third mode appeared in their *wLOD* score distribution that divided the right-hand autozygous mode in two (**Figure 3A**). While an apparent third mode also appeared in the *wLOD* score distribution of the Bengali (BEB) Asian Indian population, it was not as well defined as those of the other populations. As window size increased further, the area under both autozygous modes decreased until the left-hand autozygous mode disappeared followed sometime later by the right-hand autozygous mode. Notably, the distributions of all other populations in our dataset did not develop this third mode, and trimodality was not observed in the distribution of *LOD* scores for any of the 26 populations in the Omni2.5 dataset.

**Figure 3.**
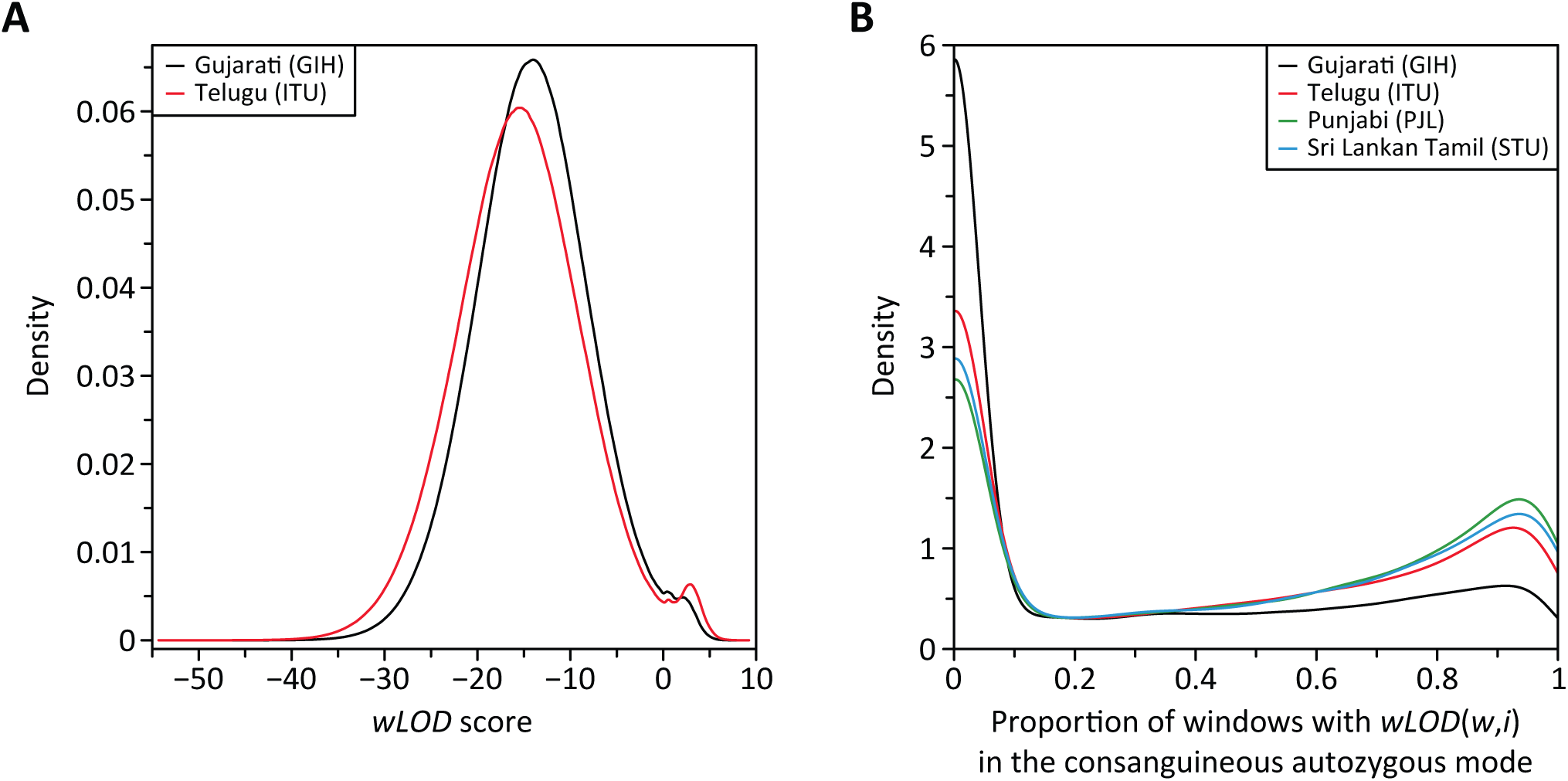
Influence of cultural processes on the distribution of scores. (**A**) Gaussian kernel density estimates of the pooled scores from all individuals in the Asian Indian Gujarati (GIH) and Telugu (ITU) populations at window sizes 200 and 220 SNPs, respectively. These patterns are representative of those observed in the Asian Indian Punjabi (PJL) and Sri Lankan Tamil (STU) populations as well as the East Asian Dai (CDX) population. (**B**) Gaussian kernel density estimates of the proportion of windows comprising each inferred ROA that are present in the right-most autozygosity mode in the Asian Indian GIH, ITU, PJL, and STU populations. ROA in the CDX population are almost exclusively in the left-most mode and it was excluded for clarity. The Asian Indian Bengali (BEB) population was excluded as we could not robustly distinguish between the two autozygous modes.

The appearance of a trimodal distribution in these six populations potentially reflects the effects of two distinct cultural processes that occur in India and among the Dai: consanguinity [172,173] and endogamy [174,175]–the restriction of marriages to within a predefined group of lineages or villages. In this scenario, the right-hand autozygous mode represents ROA due to consanguinity that are enriched for alleles rare in the general population that segregate within inbred families, while the left-hand autozygous mode represents ROA due to endogamy that are enriched for alleles present at low frequency in the general population that segregate within specific endogamic groups. Compatible with this hypothesis, the three populations with the strongest trimodal pattern (STU, ITU, and DAI) have higher reported frequencies of consanguinity (38.2% [173], 30.8% [173], and 21.3% [172]) than those with weaker trimodal patterns (BEB, 5.0% [173]; GIH, 4.9% [173]; PJL, 0.9% [173]). For example, the consanguinity-associated mode of the ITU is much larger than the endogamy-associated mode, while the reverse is true for the GIH (**Figure 3A**), consistent with consanguinity being the primary force generating ROA in the ITU while endogamy is the dominant force in the GIH. To the best of our knowledge, none of the other populations included in Phase 3 of The 1000 Genomes Project practise endogamy; consequently, we do not observe a separate endogamy-associated autozygous mode in their *wLOD* score distributions.

If trimodal distributions are indeed a reflection of the *wLOD* method being able to disentangle autozygosity signals arising from endogamy and consanguinity processes we would expect inferred ROA to be delineated predominantly by windows from only one of the two autozygous modes. Conversely, if the trimodal distribution is just an idiosyncrasy of the *wLOD* estimator we would instead expect ROA to be delineated by a random mix of windows drawn from the two autozygous modes. To investigate how windows in the putative endogamy- and consanguinity-associated modes cluster to form inferred ROA, separately for each population exhibiting a clear trimodal *wLOD* score distribution, we constructed ROA from windows with *wLOD* scores above the minimum between the non-autozygous and left-most autozygous modes in their *wLOD* score distribution [18]. Next, for each inferred ROA, we calculated the proportion of their underlying autozygous windows that had *wLOD* scores within the right-most putative consanguinity-associated mode (i.e. above the minimum between the two autozygous modes).

Inferred ROA were found to frequently be delineated by windows drawn predominantly from one of the two autozygous modes (**Figure 3B**). A large well-defined peak is observed at low proportions, representing those ROA comprised of >90% of windows drawn from the left-hand endogamy-associated mode. A more diffuse peak is observed at higher proportions, representing those ROA comprised of >80% of windows drawn from the right-hand consanguinity-associated mode. The dispersed appearance of the peak representing putative consanguinity-associated ROA can be explained as a reflection of the fact that the two autozygous modes are not distinct. At the ends of ROA arising via consanguinity, the *wLOD* scores of windows will naturally decrease as they increasingly span non-autozygous regions and overall support for autozygosity declines, leading them to increasingly fall within the endogamy-associated mode. Consequently, we would expect ROA arising via consanguinity to contain a small proportion of windows in the endogamy-associated mode, with the proportion varying based upon the overall strength of the autozygous signal (i.e. ROA conferring generally higher *wLOD* scores will have lower proportions of windows in the endogamy-associated mode). Nevertheless, across populations, 68.9% (PJL) to 84.5% (CDX) of all ROA had >80% of their component windows drawn from a single autozygous mode.

Additional support for trimodality in the *wLOD* score distribution reflecting distinct autozygosity signals arising from endogamy and consanguinity processes is provided by a comparison of how the proportion of windows drawn from the consanguinity-associated mode changes with ROA length (Additional File 1: **Figure S4**). Almost all ROA longer than 5 Mb are comprised predominantly of windows drawn from the consanguinity-associated mode (>90%), while proportions among ROA shorter than 5Mb are much more variable. This pattern is consistent with the expectation that ROA arising via consanguinity will in general be much longer than those arising via endogamy.

Overall, the properties of ROA constructed from the trimodal *wLOD* score distributions present in the Asian Indian and East Asian Dai populations are compatible with the *wLOD* method being capable of disentangling autozygosity signals that arise from different cultural processes at sufficiently large window sizes. However, further work in well-defined populations that practise both endogamy and consanguinity will be required to fully evaluate this apparent property of the *wLOD* method.

### Accuracy of the *wLOD* estimator

To evaluate the sensitivity and specificity of the *wLOD* method to detect ROA in dense genotype data, we simulated 50 independent replicates of genetic data under two demographic scenarios that are broadly representative of situations in which inbreeding and its effect on fitness are of interest as previously described [176] except that we considered a non-uniform distribution of recombination rates across the simulated chromosomes and allowed all base pairs to be mutatable (see **Methods**). Scenario 1 considered a small partially isolated population of constant effective size (*N*_e_=75) that receives approximately one migrant per generation, simulated for 150 generations (4,350 years for a generation time of 29 years [177]). Scenario 2 considered a medium sized closed population (*N*_e_=500 simulated for 100 generations [2,900 years]). Each simulated dataset consisted of a single 250 Mb chromosome upon which ∼750,000 polymorphic single-nucleotide variants (SNVs) segregate, consistent with the SNV density and length of chromosome 1 in The 1000 Genomes Project Phase 3 WGS data. The simulated WGS datasets used in downstream analyses contained 50 randomly chosen individuals from the final generation with genotypes for 709,862-746,963 SNVs in scenario 1 and 737,957-748,572 SNVs in scenario 2.

To better mimic real genetic datasets, we randomly introduced genotyping errors separately into each simulated dataset at a rate of 0.001, a conservative value that is similar to but slightly higher than the average rate of genotype discordance across 1,693 individuals between genotypes in their WGS data and those obtained at the exact same SNVs with the Illumina HumanOmni2.5 BeadChip [155], and we set ε to this value in all analyses. Analysis of the simulated pedigrees found the parents of individuals in the final generation to have a common ancestor on average three generations in the past for scenario 1 (all between 1 to 5 generations) and four generations in the past for scenario 2 (all between 1 to 7 generations) and *M* was set to these average values when analyzing their respective datasets.

Separately for each simulated dataset, we applied the *wLOD* estimator considering windows of between 50 and 500 SNPs (in 10 SNP increments), count estimates of allele frequencies calculated using all 75 individuals, and the genetic and physical map positions of each genotyped position returned by the simulation program. All windows with *wLOD* scores higher than the location of the minimum between the non-autozygous and autozygous modes in the *wLOD* score distribution were considered autozygous [18]; overlapping autozygous windows were joined to define ROA. Here, we varied the proportion of overlapping windows that must be called as autozygous when defining ROA between 0 and 50 percent (in 1% increments). As each SNV is included in multiple windows (i.e. an SNV is included in 50 different windows at a window size of 50), near the edges of a true ROA some SNV will be included in both autozygous and non-autozygous windows as the sliding window enters and leaves the ROA. Requiring an SNV to be covered by a certain proportion of autozygous windows before it is placed within an ROA can improve the accuracy of ROA inferences when using a sliding-window approach [146].

For each simulated dataset, we then calculated three measures of how well inferred ROA agreed with true ROA reported by the simulation program. First, we calculated the power of the *wLOD* method to detect true ROA, defined here as the total length of true ROA that is overlapped by inferred ROA divided by the total length of true ROA. Second, we calculated its false positive rate as the total length of inferred ROA that does not overlap with true ROA divided by the total length of inferred ROA. Finally, for all true ROA detected with the *wLOD* method, we calculated the ratio of inferred ROA length and true ROA length for all ROA. Here, ratios greater than one indicates a tendency to overcall ROA by falsely including non-autozygous regions near the boundaries of the true ROA, while ratios below one indicate a tendency to instead undercall an ROA by falsely excluding true autozygous regions near the boundaries of the true ROA [178].

As can be seen in **Figure 4A**, large numbers of false positive ROA calls are made by the *wLOD* method with a window size of 50 SNPs, decreasing markedly as the window size and the proportion of overlapping windows required during ROA construction increases. These patterns are consistent with the observation that false positive ROA calls are very small—on average 16.97 kb (standard deviation [SD] = 3.85) with a window size of 50 SNPs—and therefore delineated by a few erroneous autozygous windows that progressively fail to meet the required threshold during ROA calling as the window overlap fraction increases. Once window size reaches ∼90 SNPs, the *wLOD* estimator is able to distinguish autozygosity from homozygosity-by-chance with great precision. Conversely, numbers of false negative ROA calls increase as a function of window size and overlap fraction (**Figure 4B**). These patterns are consistent with the expectation that as window size increases smaller ROA increasingly go undetected (Additional File 1: **Figure S5A**), likely as a result of them being spanned by progressively fewer but larger windows and their autozygosity signal being increasingly masked by the inclusion of non-autozygous flanking regions in the *wLOD* score calculation. Similarly, higher overlap fractions also lead to small ROA spanned by just a small number of autozygous windows increasingly going undetected (Additional File 1: **Figure S5D**) as they fail to meet the required threshold. Nevertheless, overall power to detect ROA with the *wLOD* method only decreases slightly as window size and overlap fraction increase (**Figure 4C**), consistent with the expectation that at larger window sizes (Additional File 1: **Figure S5B**) and overlap fractions (Additional File 1: **Figure S5E**) the sliding window approach will have increasing difficulty in detecting smaller ROA but nonetheless retains high power to detect longer ROA. Finally, ratios of inferred to true ROA length increase as a function of window size and decrease as a function of overlap fraction (**Figure 4D**), reflecting the tendency of the *wLOD* method to overcall the boundaries of smaller ROA at larger window sizes (Additional File 1: **Figure S5C**) and smaller overlap fractions (Additional File 1: **Figure S5F**) with those of longer ROA affected to a much lesser extent. All together, these patterns suggest that a suitable point within the parameter space at which to evaluate the sensitivity and specificity of the *wLOD* method will be the smallest window size and overlap fraction combination at which no false-positive ROA are inferred and the average ratio of inferred to true ROA length is approximately one (**Table 3**), striking a balance between sensitivity to detect smaller ROA and the overall accuracy of ROA calls.

**Table 3.** Optimal window size and overlap proportion in the simulated datasets.

| SNV subset | Scenario 1 |  | Scenario 2 |  |
| --- | --- | --- | --- | --- |
|  | Window size | % overlap | Window size | % overlap |
| 18,000 | 60 | 7 | 70 | 5 |
| 50,000 | 70 | 18 | 80 | 19 |
| 80,000 | 80 | 28 | 90 | 22 |
| 125,000 | 80 | 29 | 100 | 26 |
| 750,0000 | 130 | 37 | 120 | 32 |

**Figure 4.**
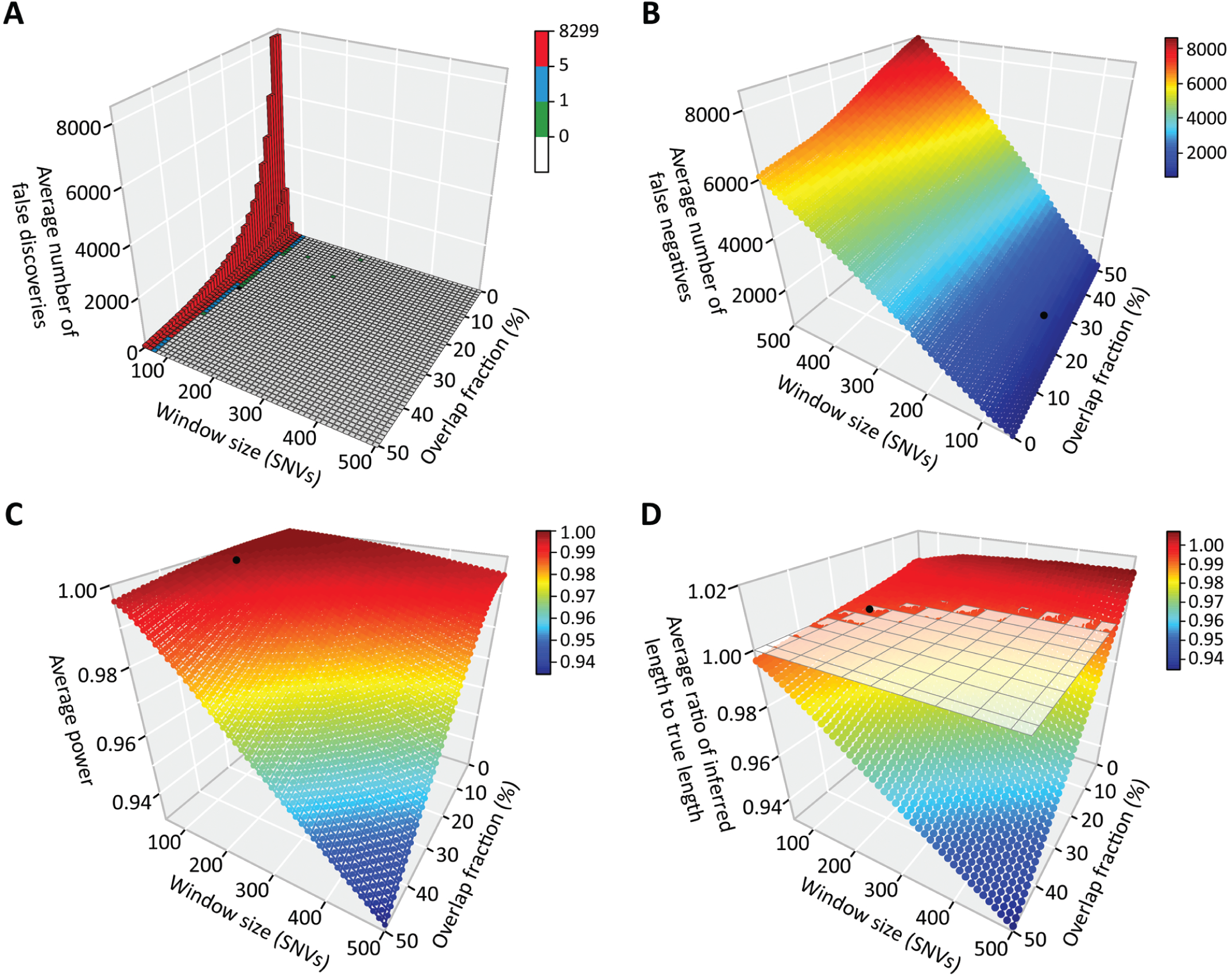
Performance of the method across different window sizes and overlap fractions. For scenario 1 and the 750,000 polymorphic SNV datasets, a three dimensional (3D) bar graph depicting the average number of falsely discovered ROA (**A**) as well as 3D scatterplots depicting the average number of false negative ROA (**B**), average power (**C**), and average ratio of inferred and true ROA lengths (**D**) reported by the method for each window size and overlap fraction across the 50 replicates are shown. In each graph, the point representing the smallest combination of window size and overlap fraction that had an average number of falsely discovered ROA of 0 and an average ratio of inferred and true ROA lengths of about 1 is shown in black.

To evaluate how SNV density influences the sensitivity and accuracy of ROA inference with the *wLOD* method we created three subsets of the simulated WGS datasets that reflect the SNV densities of commonly used human microarray-based genotyping platforms: Illumina’s HumanOmni2.5-8 (125,000 SNVs) and OmniExpress-24 (50,000 SNVs) BeadChips and Affymetrix’s Genome-Wide Human SNP 6.0 Microarray (80,000 SNVs). In addition, we included subsets with SNV densities consistent with the genotyping platforms used by ROA studies in cattle and dogs: Illumina’s Bovine HD (80,000 SNVs) and Canine HD (18,000 SNVs) BeadChips. After the removal of monomorphic SNVs, the 125K, 80K, 50K, and 18K subsets contained between 117113-123766, 74953-79211, 46846-49507, and 16865-17823 polymorphic SNVs, respectively, for scenario 1, and between 121833-122815, 77973-78602, 48733-49126, and 17544-17686 polymorphic SNVs for scenario 2. ROA were inferred and evaluated exactly as described above for the WGS datasets containing ∼750K SNVs, with the optimal window size and overlap fraction determined separately for each SNV density and demographic scenario (**Table 3**). Interestingly, optimal window size varied only slightly across the different SNV densities, lying between 60–130 SNPs and 70–120 SNPs for scenarios 1 and 2, respectively, but nevertheless increasing as a function of SNV density. The optimal window overlap fraction did however vary more widely, increasing as a function of SNV density and lying between 7–37% and 5–32% for scenarios 1 and 2, respectively.

As would be expected, the power of the *wLOD* method to detect ROA increases as a function of ROA length and the density of SNV in the genetic dataset (**Figure 5**). While ROA longer than 1 Mb are captured extremely well (>99.7%) at all SNV densities explored, the detection of ROA shorter than ∼1 Mb decreases appreciably as a function of SNV density. Nevertheless, even with only ∼18,000 SNVs (1 SNV every ∼14 kb) the *wLOD* method is able to capture 96.3% and 89.0% of ROA under scenarios 1 and 2, respectively, with this increasing to 99.9% for both scenarios with 750,000 SNVs (1 SNV every ∼333 bp). However, false discovery rates do increase dramatically with decreasing SNV density, particularly for smaller ROA (**Figure 5**) where they jump from 0.0045 and 0.0069 with 750,000 SNVs to 0.0445 and 0.1362 with 18,000 SNVs for scenarios 1 and 2, respectively, while longer ROA are much less affected: 0.0010 and 0.0001 with 750,000 SNVs increasing to 0.0200 and 0.0495 with 18,000 SNVs for ROA ≥ 5 Mb, respectively. It should be noted that these false discovery rates are solely the result of overcalling true ROA and not erroneous ROA calls. This is reflected in the ratios of inferred to true ROA length (**Figure 5**) that increase with decreasing SNV density, particularly for smaller ROA, and approach—but never quite reach—one with increasing ROA length.

**Figure 5.**
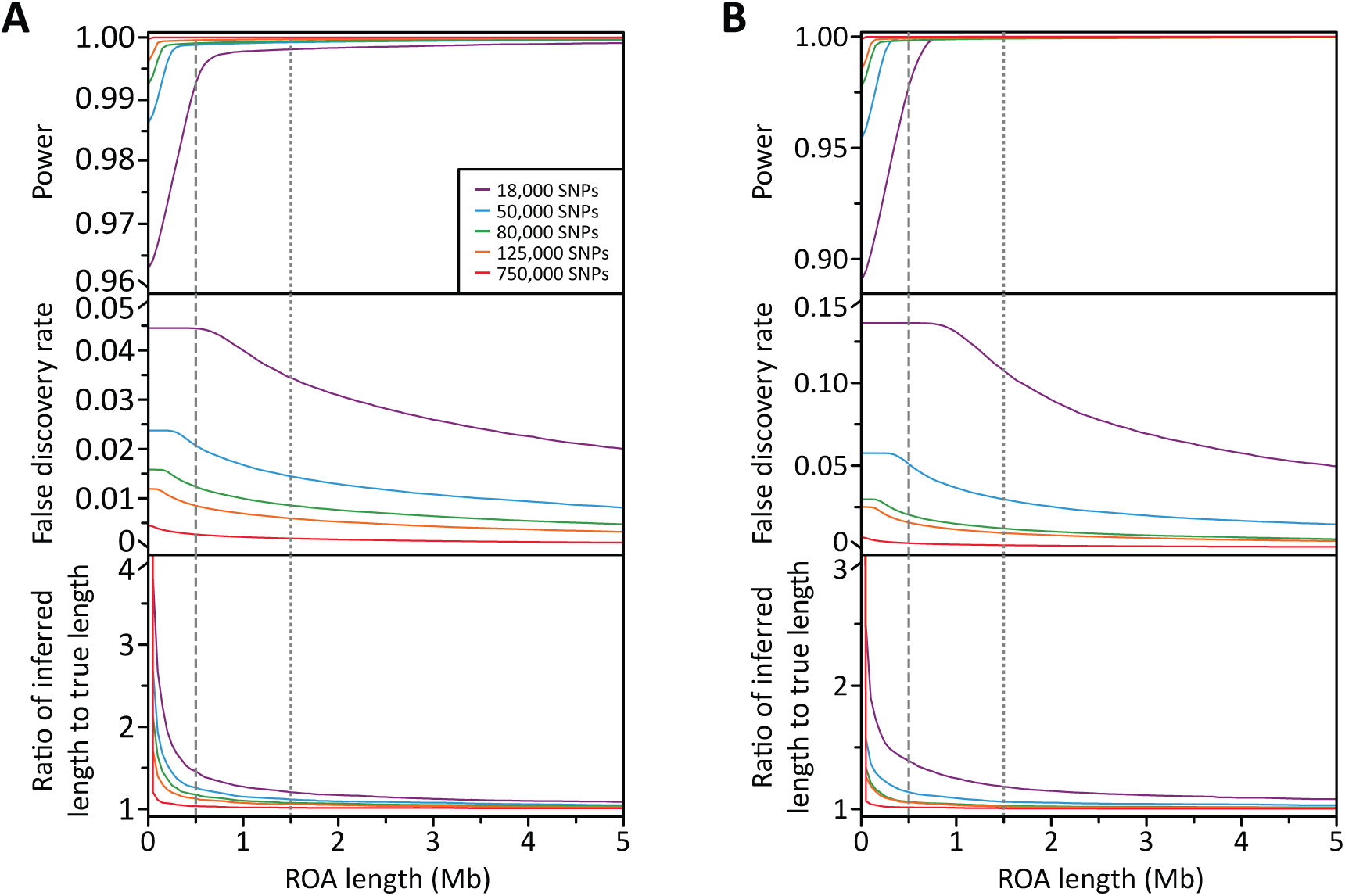
Performance of the method at different SNV densities. Lines graphs showing how average power (top), false discovery rate (middle), and ratio of inferred and true ROA length (bottom) across 50 replicate genetic simulations change with increasing ROA length for each SNV subset under (**A**) scenario 1 and (**B**) scenario 2. Each comparison was performed at the optimal combination of window size and overlap fraction for that scenario and SNV subset (**Table 3**). The grey vertical lines denote 500 kb (dashed) and 1.5 Mb (dotted), frequently applied length thresholds used to categorize ROA arising due to LD (< 500kb) and inbreeding (> 1.5 Mb) in humans [9]. Note that in scenario1, power to detect ROA >1 Mb with 18,000 SNVs surpasses that with 50-125,000 SNVs as a consequence of the optimal overlap fraction used: the overlap fraction of 0 used for the 18,000 SNV dataset is much lower than the 0.15–0.22 fractions used for the 50-125000 SNV datasets. Consequently, greater power to detect ROA >1Mb is achieved with 18,000 SNVs than is possible with 50-125,000 SNVs through less stringent placement of ROA boundaries, but at the expense of more frequent overcalling of ROA (inflated false discovery rate).

Overall, these findings indicate that the *wLOD* method is well powered to detect ROA with high sensitivity and good specificity at a wide range of SNV densities that are consistent with WGS as well as popular microarray-based platforms that are commonly used in human and non-human studies of ROA, and in particular long ROA that are of interest in studies of Mendelian and complex diseases and traits. In the simulations, both the optimal window size and the optimal overlap fraction increased logarithmically as a function of SNV density (*R*^2^=0.9814 and *R*^2^=0.8868, respectively, when considering their averages across scenarios). Fitting these averages against the natural logarithm of average SNV density *D* across all 50 replicates of their respective SNV subset, this suggests that as a rule of thumb future studies apply the *wLOD* method at a window size equal to 16.400×*log*_*e*_(*D*) + 218.020 and an overlap fraction equal to 0.0736×*log*_*e*_(*D*) + 0.8063. Based upon these equations, and calculating SNV density as the number of autosomal SNPs on the microarray divided by the total length of the target species’ autosomal genome, guideline settings for window size and overlap fraction with the commonly used human and non-human genotyping microarrays are: 111 SNPs (33%), 103 SNPs (29%), 85 SNP (21%), 81 SNPs (19%), and 59 SNP (9%) for Illumina’s HumanOmni5, HumanOmni2.5, Bovine HD, OmniExpress, and Canine HD BeadChips, respectively, and 85 SNPs (21%) for the Affymetrix Genome-Wide Human SNP 6.0 Microarray. Considering the range of autosomal SNVs observed in the WGS data available for the 26 worldwide populations in Phase 3 of The 1000 Genomes Project (12–24 million SNV [155]) a window size of 128–140 SNPs and an overlap fraction of 40–45% would be recommended for WGS datasets. Nevertheless, the modest effect window size has on power to detect longer ROA across the simulated SNV densities (Additional File 1: **Figure S6**) would suggest that the use of more conservative (i.e. larger) window sizes will not greatly impact the ability of future studies to detect longer ROA of interest regardless of the source and density of the SNV data being analyzed. The window overlap fraction used in ROA construction can then be tailored to meet the needs to detect shorter ROA (Additional File 1: **Figure S7**) and to accurately place ROA boundaries (Additional File 1: **Figure S8**), where less restrictive (i.e. smaller) fractions can greatly improve the detection of shorter ROA without significantly impacting the accuracy of longer ROA inferences.

### Performance of *wLOD* against existing ROA detection methods

We have shown the *wLOD* method to be well powered to detect ROA in genetic datasets consistent with WGS and microarray-based genotyping. We next evaluated how the power and false discovery rate of the *wLOD* method compared with those of the original *LOD* method as well as the naïve genotype counting method implemented in *PLINK* [146] and the recently reported hidden Markov model (HMM) method implemented in the *RoH* function of *BCFtools* [154] using the datasets simulated above. We do not consider here the ROA detection methods of *GERMLINE* [148] and *Beagle* [179] as they have been previously shown to underperform compared with the method implemented in *PLINK* [149]. Since the false discovery and boundary placement properties of the sliding-window-based *LOD* and *PLINK* methods would be expected to differ from those of the *wLOD* method due to their different underlying models, separately for each dataset we identified the optimal window size and overlap fraction for the *LOD* method (Additional File 2: **Table S1**) and *PLINK* (always a window size of 50 SNPs and an overlap fraction of zero) as described above. For *PLINK* we allowed at most 2% of SNPs to have heterozygous genotypes and 5% of SNPs to have missing genotypes for a window to be inferred to be autozygous [149]. The *LOD* method and *BCFtools/RoH* were applied using the same allele frequency estimates and error rate ε as the *wLOD* method, while *BCFtools/RoH* additionally incorporated genetic map positions and performed Viterbi training with initial transition probabilities between autozygous and non-autozygous states and vice versa of 6.6×10^−8^ and 5.0×10^−9^, respectively, to optimize its underlying model prior to ROA calling [154].

For both scenario 1 and 2, all four methods were able to detect >99.5% of ROA on average with 750,000 SNVs (**Figure 6A** and **6D**, respectively), representative of the density of SNVs observed in WGS data. Nevertheless, the *wLOD* method outperformed both the original *LOD* method as well as *PLINK* and *BCFtools/RoH*, particularly at shorter ROA lengths. Interestingly, power with *BCFtools/RoH* became increasingly erratic at longer ROA lengths, most noticeably in scenario 1 (small isolated populations), for reasons that remain enigmatic. However, while the *wLOD* method had a lower false discovery rate than the *LOD* method, it was notably higher than that of *BCFtools/RoH* and *PLINK*. Again, it should be noted that this elevated false positive rate solely reflects the overcalling true ROA due to the sliding-window approach employed and not erroneous ROA calls, with such overcalling easily reduced through the use of a more stringent overlap fraction but at the expense of power to detect short ROA. Nevertheless, average ratios of inferred to true ROA length were broadly similar across the w*LOD*, *LOD*, and *BCFtools/RoH* methods, where they are highest for extremely short ROA and decrease exponentially with increasing ROA length until they approach—but never quite reach—one, although ratios with *BCFtools/RoH* were marginally lower than those with the *wLOD* and *LOD* methods in scenario 2. Conversely, average ratios with *PLINK* decreased noticeably as a function of ROA length—reaching 0.47 in scenario 1 and 0.81 in scenario 2—consistent with the expectation that as a consequence of its naïve model, *PLINK* will have a tendency to undercall ROA or return fragmented ROH calls across their span as a function of the distribution of heterozygous genotypes within the ROA, which would be expected to be most numerous near its boundaries. Overall, these observations would suggest that model improvements implemented in the *wLOD* estimator (equation 2) that account for the confounding effects of LD, recombination, and mutation in the autozygosity likelihood calculation provide improved sensitivity and specificity in ROA calling over the original *LOD* estimator (equation 1). Additionally, they indicate that the *wLOD* method’s sliding window approach, which combines evidence for autozygosity across multiple SNVs, provides improved sensitivity to detect ROA compared with the HMM method of *BCFtools/RoH*, albeit with slightly decreased accuracy in ROA boundary placement.

**Figure 6.**
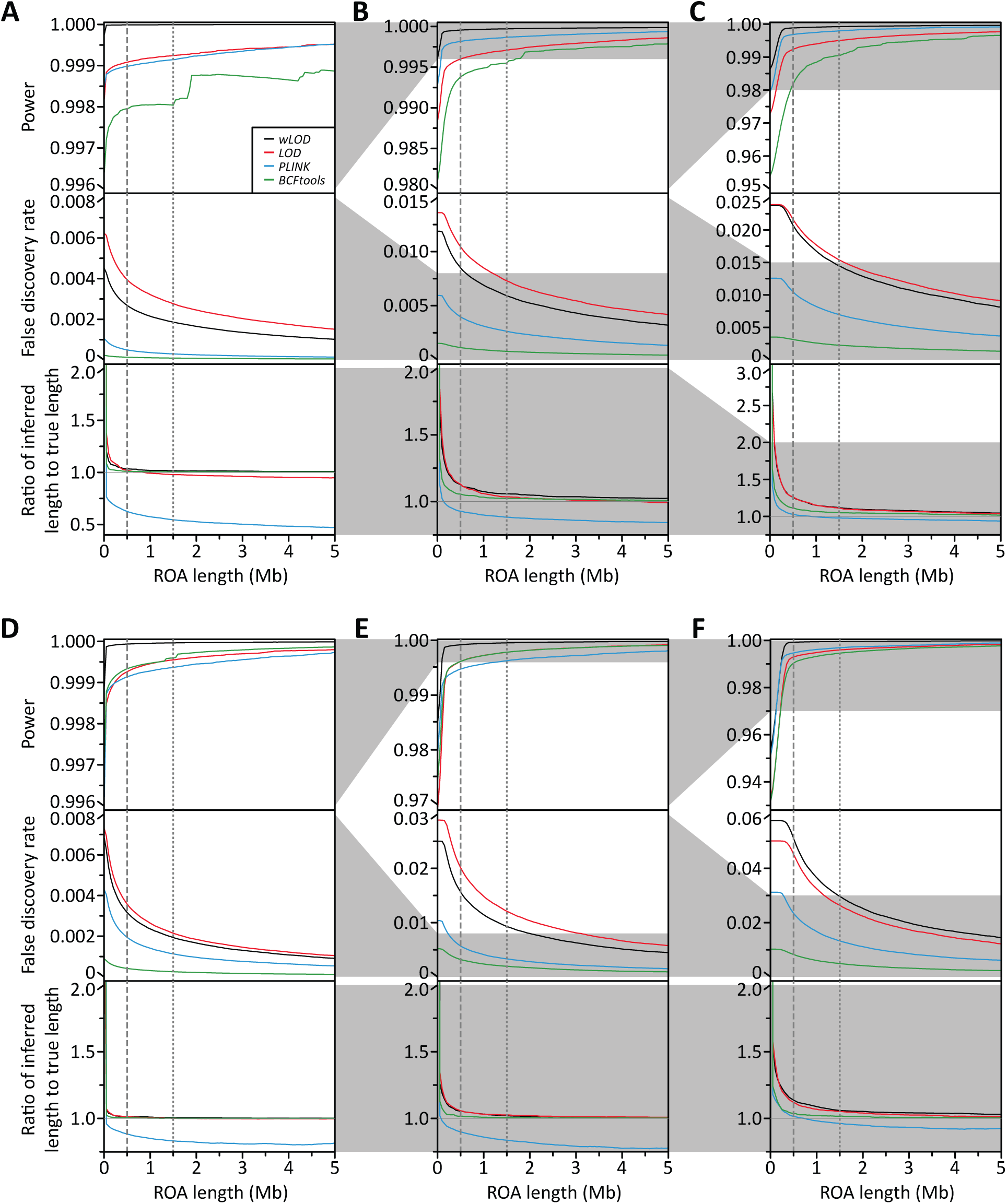
Performance of the method compared with existing methods. Line graphs showing for scenarios 1 (**A-C**) and 2 (**D-F**) and subsets consistent with WGS (750,000 SNV; **A & D**) and the Illumina HumanOmni2.5-8 (125,000 SNV; **B & E**) and HumanOmniExpress-24 (50,000 SNV; **C & F**) BeadChips how average power (top), false discovery rate (middle), and ratio of inferred and true ROA length (bottom) across 50 replicate genetic simulations change with increasing ROA length. The grey vertical lines denote 500 kb (dashed) and 1.5 Mb (dotted), frequently applied length thresholds used to categorize ROA arising due to LD (< 500kb) and inbreeding (> 1.5 Mb) in humans [9].

When we consider simulated datasets consistent with those of genotyping microarrays we observe similar patterns to those observed with 750,000 SNVs (**Figures 6** and **S9** [Additional File 1]). For both scenarios 1 and 2, the *wLOD* method consistently outperforms the *LOD* method as well as *BCFtools/RoH* and *PLINK* in terms of power, particularly at shorter ROA lengths. False discovery rates with the *wLOD* method are consistently lower than those with the *LOD* method but remain slightly higher than those with *BCFtools/RoH*, while ratios of inferred to true ROA length remain similar across the *wLOD* and *LOD* methods and *BCFtools/RoH*. As SNV density decreases from 750,000 SNVs down to 18,000 SNVs several patterns emerge. First, the difference in power between the *wLOD* and *LOD* methods decreases as a function of SNV density (Additional File 1: **Figure S9B** and **S9D**), disappearing faster under scenario 2 (large closed populations) than under scenario 1 (small partially isolated populations). These patterns are consistent with the view that in datasets containing fewer SNVs, LD confounds the inference of ROA appreciably less than in datasets containing many SNVs. Consequently, the LD correction implemented in the *wLOD* estimator (equation 2) increasing becomes less important as SNV density decreases, leading the *LOD* and *wLOD* estimators to provide broadly similar autozygosity likelihoods. Nevertheless, false discovery rates with the *wLOD* method are consistently lower than those with the *LOD* method, in agreement with the expectation that as SNV density decreases the probabilities of unobserved recombination and mutation events between genotyped SNVs increases, with the recombination and mutation corrections implemented in the *wLOD* estimator (equation 2) enabling it to better account for these events than the *LOD* estimator (equation 1). Second, ratios of inferred to true ROA length with the *PLINK* method become more similar to those of the other three methods with decreasing SNV density. This pattern is consistent with the expectation that as SNV density decreases, the number of heterozygous genotypes within ROH will also decrease, allowing *PLINK* to increasingly detect the entire ROA. Finally, the performance of *BCFtools/RoH* decreases as a function of SNV density, although an appreciable loss of power only manifests when we reach 18,000 SNVs and is more pronounced in scenario 2 than in scenario 1 (Additional File 1: **Figure S9B** and **S9D**), suggesting that its HMM is sensitive to the effects of extended LD among sparsely distributed SNVs, a situation frequently encountered in closed populations due to elevated levels of general inbreeding. It should be noted, however, that *BCFtools/RoH* was designed for next-generation whole-genome and -exome data analysis and not for sparser microarray-derived genotype datasets, so its decline in performance in such datasets is to be somewhat expected.

Contrary to expectations based on frequent discrepancies in the autozygosity status of windows with the *wLOD* and *LOD* estimators in The 1000 Genomes Project Phase 3 populations (**Figure 2**), in our simulated datasets the *wLOD* method only provided modest improvements in power and false discovery rate over the original *LOD* method (**Figures 6** and **S9**). How can we reconcile the high similarity of ROA calls with the *LOD* and *wLOD* methods in the simulated datasets with the appreciable differences in per-window autozygosity inferences made by their underlying estimators in The 1000 Genomes Project Phase 3 data? Considering the simulated datasets containing ∼125,000 SNVs, which have a comparable SNV density to that of The 1000 Genomes Project Phase 3 Omni2.5 dataset investigated in **Figure 2**, and the same window size of 150 SNPs, across the 50 replicates for scenario 1 0.519% (SD=0.496) of windows were autozygous with the *LOD* estimator but not the *wLOD* estimator, while 2.808% (SD=1.260) were autozygous with the *wLOD* estimator but not the *LOD* estimator; for scenario 2 the values were 0.153% (SD=0.169) and 5.364% (SD=1.594), respectively. While the proportion of windows autozygous with the *wLOD* estimator but not the *LOD* estimator in the simulated datasets is similar to that observed in The 1000 Genomes Project Phase 3 populations (**Figure 2D**), the proportion of windows autozygous with the *LOD* estimator but not the *wLOD* estimator is about two orders of magnitude lower than the values observed in The 1000 Genomes Project Phase 3 populations (**Figures 2C**). Thus, while we observe the expected gain in sensitivity through a reduction in the contribution of occasional heterozygotes within ROH with the *wLOD* estimator that enables improved detection of shorter ROA comprised of common haplotypes, we do not observe the expected inflation in *LOD* scores due to the confounding effects of LD among genotyped positions that leads to increased false positive ROA calls.

Based on their underlying models, we would expect the *LOD* (equation 1) and *wLOD* (equation 2) estimators to provide highly similar inferences in situations where autozygosity patterns align almost perfectly with LD patterns among genotyped SNVs and are investigated with a sufficiently high density of SNVs that the probabilities of unobserved mutation and recombination events are effectively zero. The most parsimonious explanation for the surprisingly high similarity of ROA calls made by the *LOD* and *wLOD* methods in the simulated datasets is therefore that LD patterns in these simulated datasets do not faithfully recapitulate the complexity of those found in real populations who have experienced much more complex histories than those simulated here, limiting the impact of the LD correction (equation 3) incorporated into the *wLOD* estimator. We therefore expect to observe appreciably greater improvements in the sensitivity and specificity of ROA calls with the *wLOD* method compared with the *LOD* method in real genetic data than in our simulated datasets.

### Effect of genotyped SNV density on ROA detection in real data

We have shown the *wLOD* method to be well powered to detect ROA in genetic datasets consistent with WGS and microarray-based genotyping, and to outperform a number of existing methods in terms of power although overcalling of ROA due to the sliding window approach it employs creates slightly higher rates of false discovery than a recently reported HMM model approach. While our simulations suggest that the *wLOD* method has >99.8% power to detect ROA longer than 1 Mb across SNV densities that are consistent with those frequently used in human population- and disease-genetic studies (**Figure 6**), they do not capture the diversity of historical events and sociogenetic processes that have influenced genomic autozygosity patterns in contemporary worldwide human populations. Thus, we next sought to evaluate how robust ROA inferences are among genotype datasets created via WGS and whole-exome-sequencing (WES) as well as with the popular Illumina HumanOmni2.5-8 and OmniExpress-24 BeadChips using The 1000 Genomes Project Phase 3 data.

We first developed a WGS dataset comprised of all 75,071,695 SNVs that passed our quality control criteria (see **Methods**). Next, we developed a subset of the WGS dataset that was restricted to only the 1,830,512 SNVs that are located within the genomic regions captured by the Roche Nimblegen SeqCap EZ Human Exome Library v3.0 system to mimic a whole-exome-sequencing (WES) dataset (“WES dataset” henceforth). Finally, we developed a subset of the Omni2.5 dataset that was comprised of the 676,445 SNPs that are also present on the Illumina OmniExpress-24 BeadChip (“OmniExpress dataset” henceforth). As the *wLOD* method explicitly accounts for LD among genotyped positions within a given window (equation 3) we do not consider LD pruned datasets. Similarly, since homozygosity for minor alleles at low to rare frequencies in the population is most informative for autozygosity inference with the *wLOD* estimator (Additional File 1: **Figure S1A**), we also do not consider a minor allele frequency (MAF) pruned datasets.

For the WGS, Omni2.5 and OmniExpress datasets we applied the *wLOD* method at the window size and overlap fraction suggested by our simulation analyses given their average SNV density across populations: 125 SNPs (40%), 95 SNPs (25%), and 80 SNPs (18%), respectively. As the SNV density of the WES dataset closely resembles that of the WGS dataset in the genomic regions it covers, we used the same window size and overlap fraction settings in both the WES and WGS datasets. For all datasets, μ was set to 1.18×10^−8^ [160] and *M* was set to seven, a conservative value broadly reflecting the average of effective population size estimates for populations included in The 1000 Genome Project [155,158,161]. For the Omni2.5 and OmniExpress datasets ε was set to 4.71×10^−4^, the average rate of discordance across samples between genotypes in our Omni2.5 dataset and those obtained for 1,693 of the 2,436 individuals directly with the Illumina HumanOmni2.5 BeadChip [155], while in the WGS and WES datasets ε was instead set separately for each genotype as one minus its reported likelihood. This has the potential to improve the accuracy of ROA calls in NGS datasets by incorporating the uncertainty of each genotype call into the *wLOD* score calculation, an important potential source of erroneous ROA calls in the context of their often higher and more variable per-genotype error rates compared with microarray-derived datasets [151,152]. As such, autozygous windows comprised of SNVs with low quality genotypes have a greater chance of being false-positive signals than those with higher quality genotypes, while low quality heterozygous genotypes—that in one possibility may be genotype calling errors—located in runs of higher quality homozygous genotypes have the potential to mask true autozygous signals.

For each dataset and population, we defined a *wLOD* score autozygosity threshold as the location of the minimum between the non-autozygous and autozygous modes in its *wLOD* score distribution [18]. Sample size was not observed to appreciably influence the location of the minimum between the non-autozygous and autozygous modes (Additional File 1: **Figure S10**). However, across 100 random samples of individuals greater consistency in its determination was observed with increasing sample size, particularly compared with sample sizes of less than 10 individuals, indicating that 10 or more individuals should be used to ensure a robust estimate of the threshold is obtained. All windows with *wLOD* scores above threshold were considered autozygous [18], and overlapping autozygous windows were joined to define ROA contingent on the window overlap fraction used for that dataset.

Comparing ROA identified in the WGS and Omni2.5 datasets, we find Omni2.5 ROA to be frequently longer than their corresponding WGS ROA and in most cases to completely encompass the WGS ROA (**Figure 7A**). The magnitude of their length discrepancies decreases with increasing ROA length, consistent with the expected effects of decreased SNV density on the accuracy of inferred ROA boundaries. In addition, while all Omni2.5 ROA are present in the set of WGS ROA, the reverse is not true (**Figure 7B**). Many short ROA (<500 kb) detected in the WGS dataset are not found in the Omni2.5 dataset, with the fraction of missing ROA decreasing with increasing distance from Africa, reflecting the effect of increasing LD [162,163] on our ability to detect shorter ROA with the sparser set of SNVs in the Omni2.5 dataset. Concordance between the WGS and Omni2.5 datasets for intermediate (500 kb to 1.5 Mb) and long (>1.5 Mb) ROA is generally high, although in many populations the fraction of WGS ROA missing in the set of Omni2.5 ROA remains nontrivial. These fractions generally increase as a function of distance from Africa, likely reflecting the reduction in haplotype diversity with decreasing genetic diversity [95,164–167] decreasing our ability to distinguish autozygosity from homozygosity-by-chance, particularly over extended genomic regions when genotypes are only available for a fixed set of SNVs that were selected for their generally high level of polymorphism worldwide.

**Figure 7.**
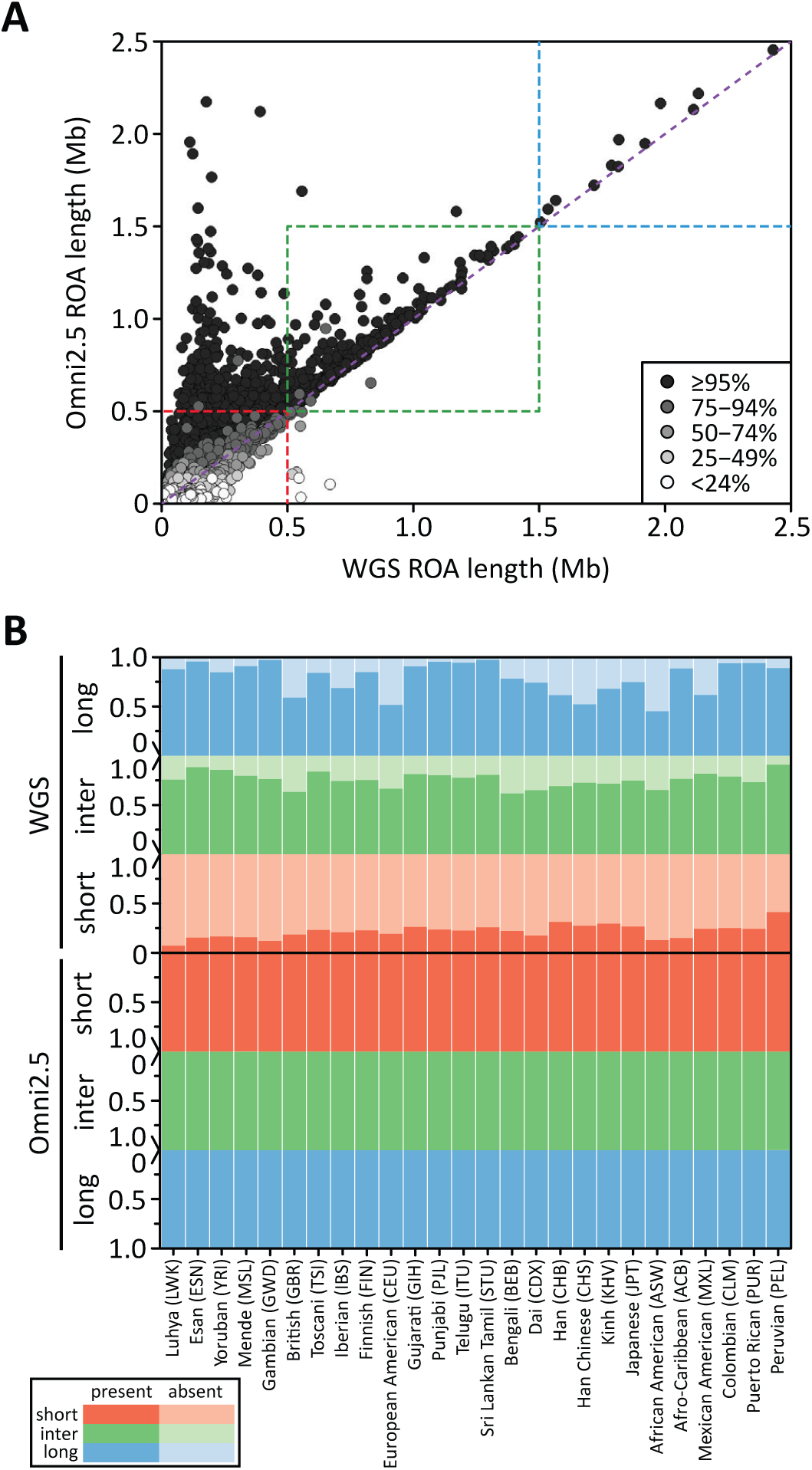
Concordance of ROA inferred in the WGS and Omni2.5 datasets. (**A**) A scatterplot comparing the length of each WGS ROA with that of its corresponding Omni2.5 ROA in the European American (CEU) population. Each point is shaded according to the proportion of the WGS ROA that overlaps the Omni2.5 ROA. (**B**) Bar plots representing the proportions of short (<500 kb; shown in red), intermediate (500 kb to 1.5 Mb; shown in green), and long (>1.5 Mb; shown in blue) ROA in the WGS (upper) and Omni2.5 (lower) datasets that overlap (darkest shade) or are absent from (lightest shade) the other dataset in each population.

Similar patterns are observed when we compare ROA identified in the Omni2.5 and OmniExp datasets, where almost all OmniExp ROA are present in the set of Omni2.5 ROA (Additional File 1: **Figure S11B**) and encompass their generally shorter corresponding Omni2.5 ROA (Additional File 1: **Figure S11A**). While many short ROA detected in the Omni2.5 dataset are not found in the OmniExp dataset, both intermediate and long ROA are captured extremely consistently between the two datasets despite their different SNV densities. Likewise, when we compare ROA identified in the WGS and WES datasets, almost all WES ROA are present in the set of WGS ROA (Additional File 1: **Figure S12B**) and tend to encompass their generally shorter corresponding WGS ROA (Additional File 1: **Figure S12A**). However, while numbers of short and intermediate ROA identified in the WGS dataset but not the WES dataset are much higher than in the same comparison between the WGS and Omni2.5 datasets (**Figure 7**), the numbers of long ROA identified in the WGS dataset but not the WES dataset are instead similar. This indicates that the non-uniform and often sparse distribution of SNVs in the WES dataset does not impact the detection of long ROA more than would be expected following a general reduction in SNV density.

Overall, these findings are consistent with the higher density of SNVs in the WGS dataset and the presence of many more rare and low-frequency SNVs detected by NGS compared with microarray-based genotyping platforms—which are particularly informative about autozygosity under our likelihood model (Additional File 1: **Figure S1A**)—greatly improving our ability to detect ROA. Nevertheless, many long ROA that are of interest in Mendelian and complex disease studies are well captured by the sets of SNVs included on Illumina’s HumanOmni2.5-8 and OmniExpress-24 BeadChips. However, the sparse and non-uniform genomic distribution of SNVs in the WES dataset creates difficulties when inferring short and intermediate ROA with the *wLOD* method, despite the presence of rare and low-frequency SNVs, while long ROA are instead captured almost as well as with genotyping microarrays. We therefore do not recommend using the *wLOD* method to detect ROA in WES datasets generated by future studies.

### Classification of ROA

ROA of different lengths reflect homozygosity for haplotypes inherited IBD from common ancestors at different depths in an individual’s genealogy: longer ROA most likely arise due to recent ancestors and shorter ROA due to more distant ancestors. We previously advocated that ROA be classified into *G* length-based classes using the Gaussian mixture model approach applied on their physical map lengths (in bp) that groups ROA based upon their supposed ages [18]: (A) short ROA that measure tens of kilobases and that are of the length at which baseline patterns in LD in a population produce autozygosity through the pairing of two copies of the same ancient haplotype, (B) intermediate length ROA that measure hundreds of kilobases to several Mb and that are likely the result of background relatedness–recent but unknown kinship between parents due to limited effective population sizes–and (C) long ROA that measure multiple megabases and are likely the result of recent parental relatedness. The choice of *G* = 3 was motivated by the observation that at *G* > 3, the additional classes were not discrete; that is, they were encompassed by one of the existing classes (Additional File 1: **Figures S13A** and **S13C**).

This classification approach is limited by the imperfect correlation between physical map lengths and genetic map lengths (Additional File 1: **Figure S14**), a more accurate representation of the relationship between ROA length and age [180,181] that is not biased by the non-uniform genomic distribution of recombination rates [182]. If we instead classify ROA based on their genetic map length (in cM) using a Gaussian mixture model we find that regardless of the number of classes considered they are always discrete (Additional File 1: **Figures S13B** and **S13D**). This would suggest that the original loss of discreteness when classifying based upon physical map length may reflect the confounding effects of physically long but genetically short (and vice versa) ROA on the overall length distribution. Nevertheless, regardless of whether physical or genetic map lengths are used the overall pattern of fit with increasing class number remains highly similar (Additional File 1: **Figures S13A** and **S13B**, respectively), where Bayesian Information Criterion (BIC) likelihoods plateau at around *G* = 5 with the WGS and Omni2.5 data and at around *G* = 4 classes with the OmniExpress data (not shown). The smaller class number for the OmniExpress dataset compared with the WGS and Omni2.5 datasets is consistent with the expectation that smaller ROA will be poorly captured by its sparser set of SNVs, ultimately leading to the loss of the shortest ROA class detected in the WGS and Omni2.5 datasets. Note that for all populations the maximum BIC likelihood is reached at *G* > 5. Future studies investigating fine scale ROA patterns may wish to consider values of *G* at which BIC is maximized, however for illustrative purposes we consider *G* = 5 here since the increase in BIC at *G* > 5 is small.

When considering a five class classification scheme, the longest class (*G* = 5) contains ROA that likely arise from recent parental relatedness and the penultimate longest class (*G* = 4) contains ROA that likely arise from recent population processes, while the shortest classes (*G* = 1-3) contain ROA arising through the pairing of two copies of much older haplotypes that have common ancestors at different times in the distant past. Sample size was observed to have a greater effect on ROA classification (Additional File 1: **Figure S15**) than on *wLOD* score threshold (Additional File 1: **Figure S10**), with the proportion of ROA whose classification differed from that assigned when all available individuals are used decreasing as a function of sample size. Importantly, the proportion of misclassified ROA decreases with increasing ROA class, with those in the longest class (*G* = 5) infrequently misclassified (mean=0.052 with SD=0.029 across all 26 populations at a sample size of 25) while those in shorter classes were more frequently affected (mean=0.092 with SD=0.046, mean=0.091 with SD=0.045, mean=0.083 with SD=0.045, and mean=0.068 with SD=0.042, for *G* = 4 to 1, respectively). These observations indicate that sample size is an important factor when classifying ROA using a Gaussian mixture model, but in general samples sizes of at least 25 individuals should provide reasonably robust classification of ROA using this approach, particularly longer ROA that are of interest in genetic studies on Mendelian and complex diseases.

### Geographic patterns in ROA

We have shown the *wLOD* method to be well powered to detect ROA in genetic datasets consistent with WGS and microarray-based genotyping, while our investigation of a Gaussian mixture model approach for ROA classification based upon their genetic map lengths indicates the presence of five ROA classes in The 1000 Genomes Project Phase 3 populations, a higher number than was used in our earlier study of the Human Genome Diversity Panel (HGDP) and International HapMap Project (Phase 3) populations that used a microarray-derived dataset and classified ROA based upon their physical map lengths [18]. To evaluate how genome-wide patterns in ROA inferred with the *wLOD* method and classified into five classes via a Gaussian mixture model applied to their genetic map lengths compared with those of earlier studies, we performed the first high-resolution survey of ROA patterns in The 1000 Genomes Project Phase 3 populations based upon ROA inferred in the WGS dataset as described above.

Consistent with previous studies [12,18,22], ROA of different lengths have different continental patterns among the 26 populations included in Phase 3 of The 1000 Genomes Project both with regards to their total lengths (**Figure 8**) in individual genomes as well as in their non-uniform distributions across the genome (**Figure 9**) that are correlated with spatially variable genomic properties such as recombination rate (Additional File 1: **Figure S16**) and signals of natural selection (Additional File 1: **Figure S17**), reflecting the distinct forces generating ROA of different lengths. Total lengths and numbers of ROA in the shortest (*G* = 1-3) and to some extent intermediate (*G* = 4) classes increase with distance from Africa, rising in a stepwise fashion in successive continental groups (**Figures 8**), in agreement with the observed reduction in haplotype diversity with increasing distance from Africa [162,183–185]. Those of the longest class (*G* = 5) do not show a similar stepwise pattern, instead exhibiting higher and more variable values in populations where consanguinity in more frequent (**Table 2**) and inbreeding coefficient estimates are generally higher [186]. Notably, the East Asian Dai have remarkably high total lengths of short ROA (G = 1-3), potentially reflecting their small population size—∼1.2 million in Yunnan province, China [187], where The 1000 Genomes Project samples were collected—and complex evolutionary history [188,189].

**Figure 9.**
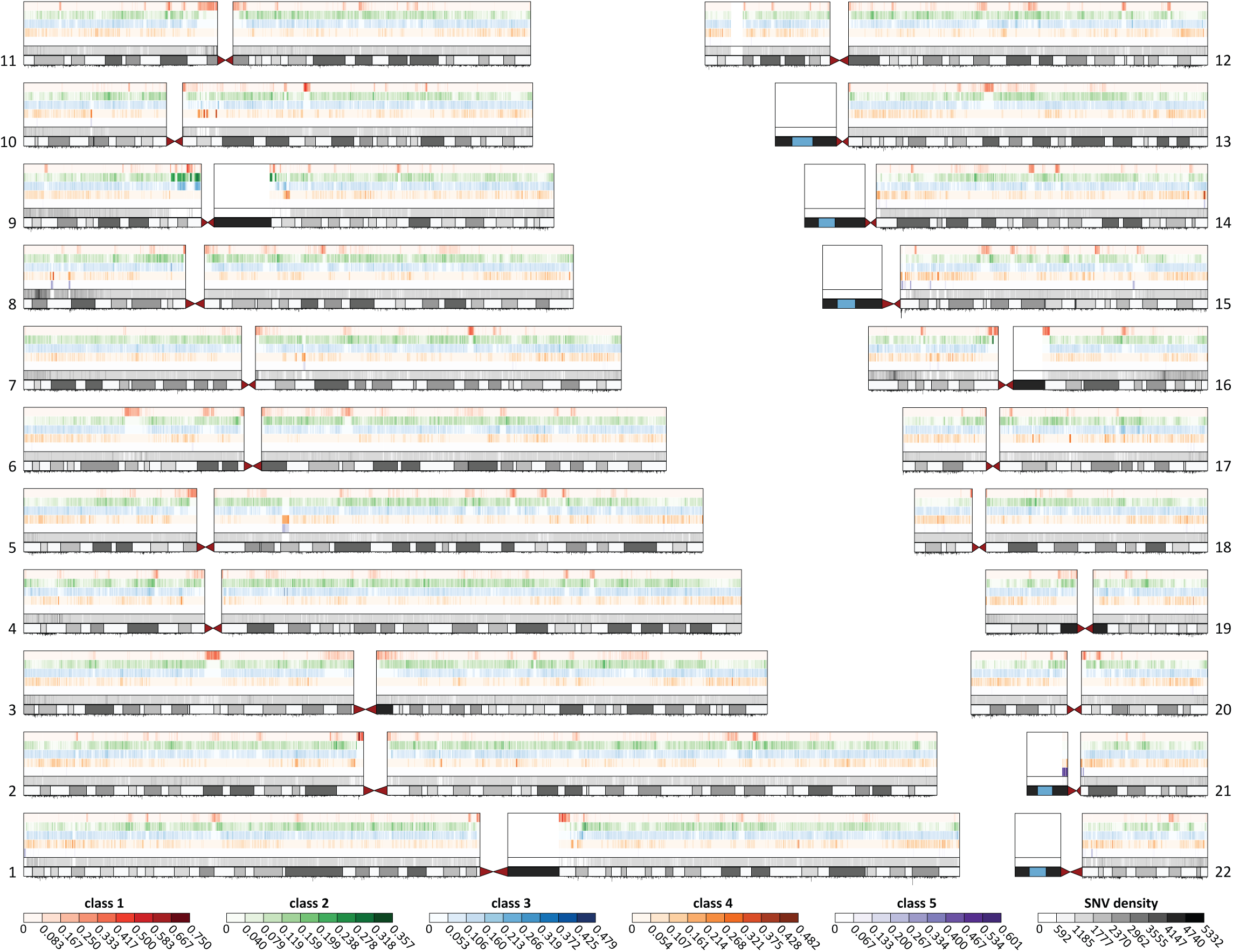
Distribution of worldwide ROA frequencies across the genome. For each autosome, the figure shows for each ROA length class the average proportion of individuals in the WGS dataset who have an ROA overlapping SNVs within non-overlapping 50 kb windows. Each row represents an ROA class, and each column represents a window. The intensity of a point increases with increasing average ROA frequency, as indicated by the color scale below the figure. The SNV density of each window and an ideogram of chromosome banding are shown in the bottom tracks, with average recombination rate in each window represented by a vertical black line below the ideogram, where line heights proportional to average recombination rate.

**Figure 8.**
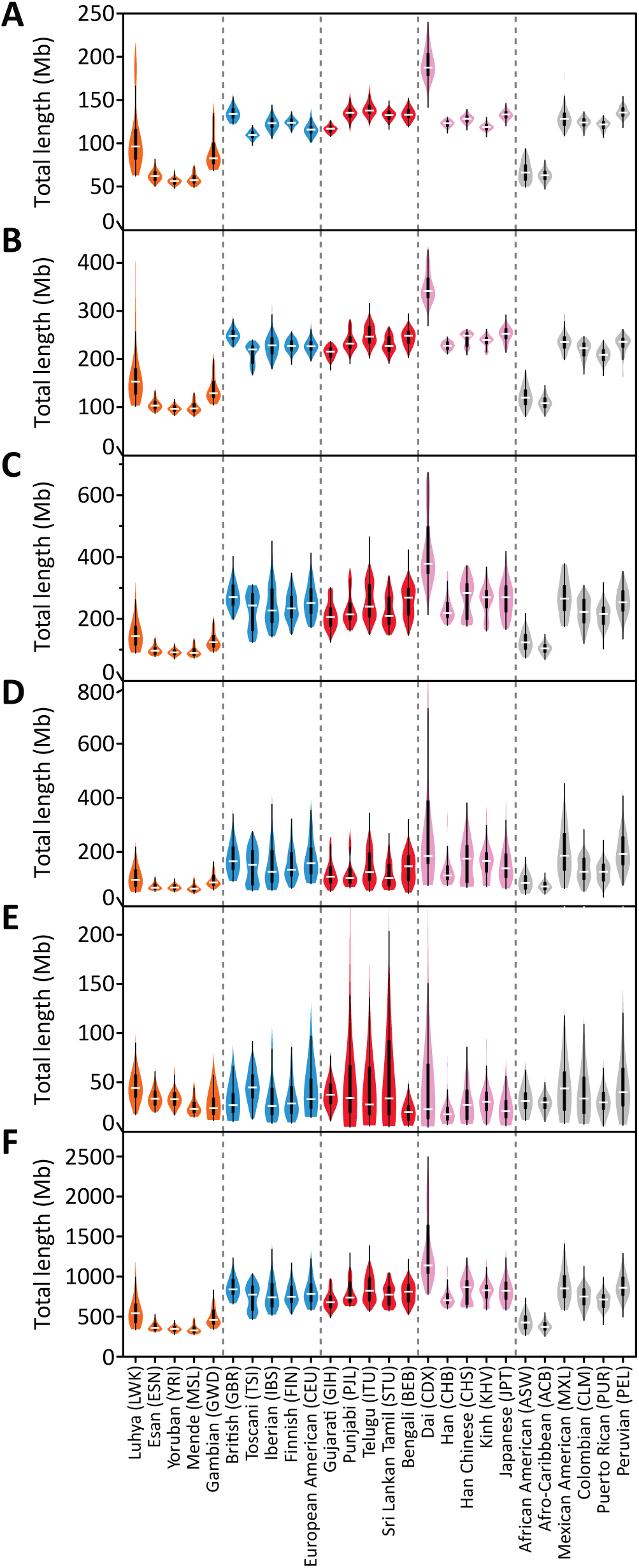
Population-specific distributions of the total length of ROA per individual. Data are shown as violin plots [249], representing the distribution of total ROA length across all individuals in each of the 26 populations for (**A**) class 1, (**B**) class 2, (**C**) class 3, (**D**) class 4, (**E**) class 5, and (**F**) all five ROA classes combined. Populations are ordered from left to right by geographic region and within each region by increasing geographic distance from Addis Ababa.

#### Recombination and natural selection

The strength of the correlation between the genomic distribution of ROA and recombination rate decreases with increasing ROA class (Additional File 1: **Figure S16**), consistent with the expectation that the patterns of genetically shorter ROA will be determined by recombination to a greater extent than longer ROA, which due to their more recent origins have had fewer opportunities for recombination events to systematically influence their patterns. Conversely, the correlation between ROA patterns and signatures of natural selection is strongest for class 2-3 ROA, and to some extent intermediate class 4 ROA, while it is very weak for the shortest (*G* = 1) and longest (*G* = 5) ROA classes (Additional File 1: **Figure S17**). These patterns are compatible with natural selection having primarily influenced genomic diversity patterns in the distant past, with autozygosity for the relics of the haplotypes that arose during those events manifesting as class 1-4 ROA, dependent upon how long ago the event occurred.

The long term effects of natural selection on patterns of ROA might be expected to be most evident in genomic regions encompassing genes implicated in one or more Mendelian diseases, where purifying selection acting on strongly deleterious alleles, which may occur more frequently in such genes due to their apparent importance for human health, would be expected to increase levels of homozygosity relative to genes much less frequently subjected to purifying selection. Using the union of two previously reported lists of genes associated with autosomal dominant (669) and recessive (1130) diseases in the Online Mendelian Inheritance of Man (OMIM) database [190–192], we created a list containing genes not associated with autosomal dominant or recessive diseases (24,260; “non-OMIM” henceforth); genes associated with both autosomal dominant and recessive diseases were ignored. For each individual, we then calculated the fraction of the total lengths of all autosomal dominant, autosomal recessive, or non-OMIM transcribed regions that are overlapped by ROA based on their genomic positions in build HG19 of the University of California – Santa Cruz (UCSC) reference genome assembly. Strikingly, regardless of the ROA length class considered, the fraction for OMIM dominant genes was almost always higher than that of non-OMIM genes (*P*<10^−16^ in all comparisons; Wilcoxon signed rank test), while the opposite was true for OMIM recessive genes (*P*<10^−16^ in all comparisons; Additional File 1: **Figure S18**). Nevertheless, the pattern is strongest for intermediate length ROA classes (*G* = 2-4) and weakest for the shortest (*G* = 1) and longest (*G* = 5) classes. Together, these results are compatible with deleterious alleles occurring less frequently in non-OMIM genes than in OMIM dominant genes, where they are efficiently removed from the population via purifying selection acting on both their homozygous and heterozygous forms, creating increased autozygosity at lengths consistent with population-level processes rather than inbreeding. One possible explanation for the decreased autozygosity around OMIM recessive genes compared with non-OMIM genes would be increased embryonic lethality and/or childhood mortality with individuals homozygous for deleterious recessive mutations in OMIM recessive genes, leading to reduced autozygosity in genomic regions encompassing them in the extant population.

Genes that have been the target of positive selection might be expected to reside within genomic regions that are more frequently autozygous in the general population than those harboring genes that have not. Considering the fraction of each gene’s transcribed region that is in a ROA in each individual’s genome, we compared their median fraction across individuals in each population (Additional File 1: **Figure S19**). While most genes have a median fraction of about zero, a number of genes that lie within genomic regions spanned by ROA in more than 90% of individuals in a population. Across populations, we observe 54 such instances with long class 5 ROA that represent seven distinct genomic regions (Additional File 2: **Table S2**), 159 with intermediate length class 4 ROA (22 distinct regions; Additional File 2: **Table S3**), and 31 (nine distinct regions; Additional File 2: **Table S4**), seven (five distinct regions; Additional File 2: **Table S5**), and 480 (46 distinct regions; Additional File 2: **Table S6**) with short class 1–3 ROA, respectively. While most genes in these regions fall within the non-OMIM group, two of the genes enriched for class 4 ROA (*CFC1* and *SMN1*) and nine of the genes enriched for class 1 ROA (*SLC25A20*, *NDUFAF3*, *LAMB2*, *GPX1*, *NPRL2*, *ACY1*, *MRPS16*, *LCAT*, and *COX4I2*) are from the OMIM recessive group, while one gene enriched for class 1 ROA is from the OMIM dominant group (*THAP1*). Future investigation of genes that are unusually frequently overlapped by ROA in the general population may provide new insights into the role of recessive variation in human phenotypic diversity and common disease risk as well as the genes within which such variation acts.

#### Genomic distribution

Genomic distributions of shorter ROA (*G* = 1-4) are similar among populations from the same geographic region (Additional File 1: **Figures S20B-E**) and closely mirror the patterns of pairwise *F*_ST_ among populations (Additional File 1: **Figure S20A**; Procrustes similarity statistic *t*_*0*_>0.803), while those of the longest ROA class (*G* = 5) vary more widely among populations (Additional File 1: **Figure S20F**; *t*_*0*_=0.466). Overall, these patterns are consistent with the interpretation that shorter ROA (*G* = 1-4), for which neighboring populations have similar patterns, reflect autozygosity that arises through population processes on different evolutionary timescales, while longer ROA (*G* = 5), for which neighboring populations do not necessarily have similar patterns, reflect autozygosity that instead arises through more recent cultural processes such as inbreeding [18].

#### Autozygosity hotspots

The non-uniform genomic distribution of the different ROA classes and their variability among populations creates autozygosity hotspots that are in some instances shared among subsets of the populations. For example, there is a hotspot for class 4 ROA on the q-arm of chromosome 2 that is common to three of the five European populations and encompasses the human lactase gene (*LCT*; **Figure 10**) that was not detected in our original study of the HGDP and HapMap populations that included 10 from Europe [18]. In this genomic region, we observe high frequencies of intermediate length class 4 ROA in the Northern European FIN and GBR populations as well as the European American (CEU) group, but not in the Southern European TSI and IBS populations or any other population in the dataset. The presence and absence of this hotspot broadly reflects worldwide patterns in lactase persistence frequency [193,194]. Lactase persistence is most frequent in Northwestern Europe [195,196] where it is caused primarily by a single mutation in *LCT* that rose to high frequency as a consequence of natural selection in response to the rise of milk consumption and pastoralism [194,197,198]. It decreases in frequency through Eastern and Southern Europe and Central/South Asia reaching near-zero frequencies in East Asia and the Americas [193,196,199–201], while it is present to varying degrees in admixed Mestizo [202–204] and African American [199,204] populations as a consequence of their recent European ancestry. Thus, we observe high levels of autozygosity around *LCT* in the GBR, FIN, and CEU populations and markedly lower but noticeable levels in the IBS, but no observable signal in the TSI or any of the Asian or admixed populations. While lactase persistence is present at moderately high frequency in sub-Saharan Africa it is caused by several different mutations [194,205] and the African populations included in The 1000 Genomes Project are located predominantly in historically non-milking areas of the continent [197]. Consequently, we do not observe a similar autozygosity signal in the African populations as we do in the Northern European populations.

**Figure 10.**
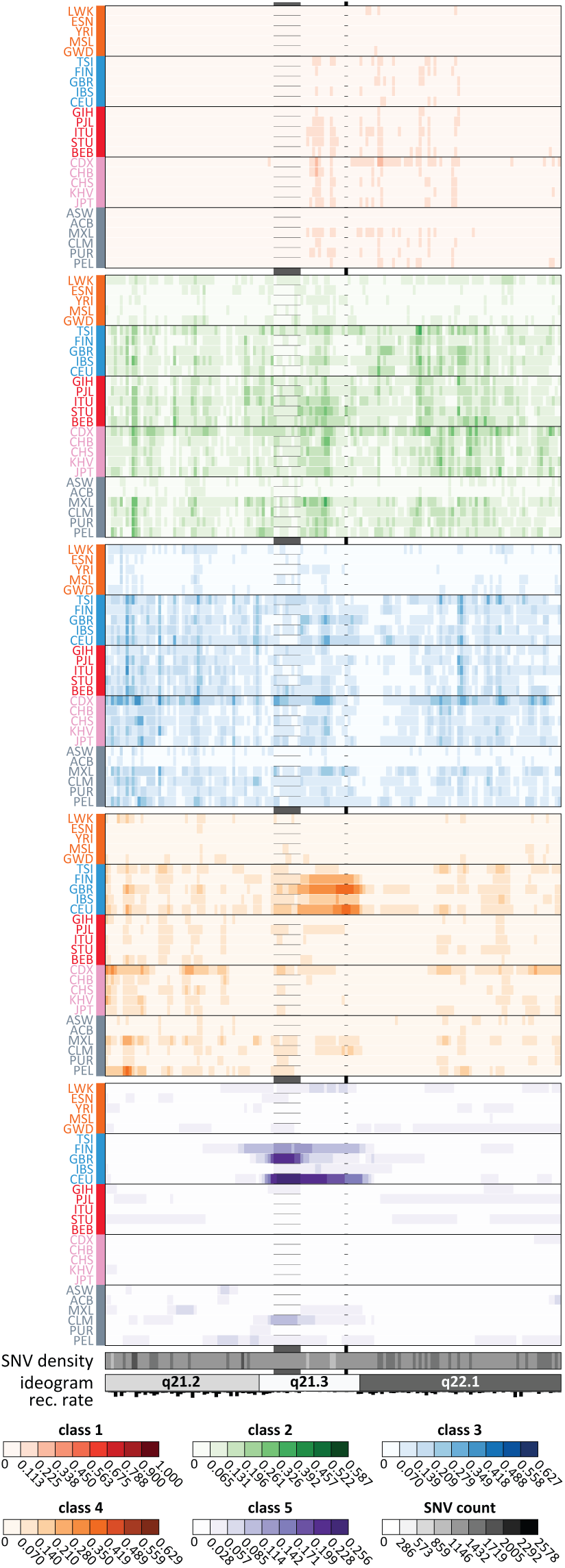
Per-population ROA frequencies within a ROA hotspot on chromosome 2. For each ROA class, for each population, the average proportion of individuals in that population who have an ROA overlapping SNVs within non-overlapping 50 kb windows from 132,500,000 to 140,200,000 bp on the q-arm of chromosome 2 is shown. Each row represents a population, and each column represents a window. Populations are ordered from top to bottom by geographic affiliation, as indicated by the color of their label, and within regions from top to bottom by increasing geographic distance from Addis Ababa (in the same order as in **Figure 8**). Average ROA frequency, average SNV density, chromosome banding, and recombination rates are shown as in **Figure 9**. The black vertical box demarks the location of the *LCT* gene, while the vertical grey box demarks the location of the class 5 ROA hotspot in the CEU and GBR.

Interestingly, we also observe a hotspot for the longest ROA class (*G*=5) at the same location in the Northern European CEU and GBR populations ∼770kb downstream of the *LCT* gene (**Figure 10**), while a weaker spike in class 5 ROA frequency is seen in the FIN population. This hotspot encompasses four genes within its core region (chr2:135,375,000-135,775,000) that encode a transmembrane protein (*TMEM163*), an aminocarboxymuconate semialdehyde decarboxylase (*ACMSD*), cyclin T2 (*CCNT2*), and a mitogen-activated protein kinase kinase kinase (*MAP3K19*). The maximum normalized haplotype-based selection statistic *nS*_L_ [206] score observed in the CEU, GBR, and FIN populations within the core region is 4.980, 4.818, and 4.962, respectively, suggesting that this ROA hotspot potentially reflects the outcome of recent positive selection. However, none of the genes within this hotspot are known to have functional consequences when mutated, leaving the cause of this ROA hotspot and its putative signals of positive selection enigmatic.

Overall, frequency patterns in this genomic region of the different ROA classes in the Northern European CEU, GBR, and FIN populations are consistent with positive selection having occurred at two different time-points. The extended haplotypes created by historical positive selection acting on the single *LCT* mutation that arose in ancestral Northern Europeans have, over subsequent generations, decreased appreciably in length, but due to the marked reduction in haplotype diversity in the surrounding region commonly create intermediate length class 4 ROA through background population processes. Conversely, the presence of extended IBD haplotypes creating longer class 5 ROA in a genomic region ∼770 kb away from *LCT* would be compatible with positive selection acting much more recently, in agreement with the atypically high *nS*_L_ scores observed within this region in these populations.

### Statistical inference of enrichment of autozygosity signals between groups

A unique feature of the *wLOD* ROA detection approach is the availability of log-likelihoods of autozygosity for each window in each individual examined. It is therefore possible to directly compare the strength of autozygosity signals between two or more groups of individuals to identify those windows that have significantly greater evidence for shared autozygosity signals in one group compared with the others [150]. In one possibility, such an approach could be used to identify genomic regions that have stronger signals of autozygosity in affected versus unaffected individuals and thus may harbor disease-associated mutations. Similarly, genomic regions with significantly stronger signals of autozygosity in one subset of a population compared to another other may reflect founder effects if there is limited gene flow between them or the presence of adaptive alleles in one subset but not the other that have risen to high frequency.

We demonstrate the principle of this approach using three of the five Central/South Asian groups included in Phase 3 of The 1000 Genomes Project who represent subpopulations within the larger Indian population: BEB, GIH, ITU, PJL, and STU. Genetic diversity patterns in these five groups support the presence of two genetically distinguishable clusters within the GIH, ITU, and PJL (Additional File 1: **Figure S21**). When instead compared pairwise, the larger of the two ITU clusters lies intermediate between the smaller ITU cluster and the larger of the two GIH, PJL, or STU clusters, while the largest of the PJL clusters overlaps significantly with the smaller GIH cluster (not shown). The GIH individuals were sampled in Houston, TX, while the BEB, ITU, PJL, and STU individuals were all sampled in the UK. Given the intermediate locations of the larger ITU and PJL clusters in the pairwise comparisons, they may potentially reflect admixed individuals within these sample sets. However, both clusters are tightly bunched arguing against this possibility given the normal dispersion of admixed individuals in such analyses owing to their continuum of admixture levels [207,208]. In another possibility, these distinct clusters might represent the unintentional sampling of distinct endogamic communities whose restrictive marital practices under the long-established Indian caste system has made them distinguishable genetically [209].

Because we would expect differential autozygosity signals among groups to have arisen relatively recently through population or cultural processes, window size is not constrained by our power to detect shorter, more ancient, ROA. A natural window size to use when searching for differential autozygosity signals between groups is therefore the one whose *wLOD* score distribution can best discriminate between autozygous and non-autozygous windows. In one possibility, this can be defined as the window size that maximizes the distance between the autozygous and non-autozygous modes—measured here as the distance between the modal score in each mode (**Figures 1B** and **S2** [Additional File 1]). Using the WGS dataset and optimal window sizes of 450, 580, and 610 SNVs for the GIH, PJL, and ITU, respectively, we compared the *wLOD* scores of individuals present in each of their two clusters (Additional File 1: **Figure S21**) and evaluated the significance of their observed differences with the permutation-based approach described in Wang *et al*. [150] except that here we use a Wilcoxon rank-sum test instead of the two sample *t*-test suggested by Wang *et al*. as it is much less sensitive to the presence of outliers but has similar power to detect a location shift [210]. Briefly, separately for each group, we first create a distribution of test statistics under the null hypothesis of no difference in *wLOD* scores between clusters using 1,000 permutations of cluster labels, recording for each permutation the maximum observed test statistic across all windows genome-wide. Next, separately for each window, a genome-wide adjusted *P*-value for the significance of the observed differences in *wLOD* scores between clusters is then calculated as the proportion of the maximum genome-wide test statistics observed in the 1,000 permutations that exceeded the test statistic obtained with the true labels for that window. Finally, for each cluster, genomic regions enriched for autozygosity signals in that cluster compared with the other were defined by joining together overlapping windows with a permutation *P*-value (*P*_perm_) ≤ 0.05.

Intriguingly, while we would not *a priori* expect to observe significant differences in the strength of autozygosity signals between the two apparent clusters within the GIH, ITU, and PJL sample sets, we did identify one genomic region significantly enriched for autozygosity signals in cluster A compared with cluster B in both the ITU and PJL (**Figures 11B** and **11C**; **Table 4**); no regions were identified in the GIH (**Figure 11A**). The genomic region in the ITU lies within the transcription elongation regulator 1 like (*TCERG1L*) gene that has been associated with regulation of plasma levels of the adipokine adiponectin [211], a modulator of glucose regulation and fatty acid oxidation [212] implicated in obesity, diabetes, coronary artery disease and Crohn’s disease risk [213–215]. The genomic region in the PJL encompasses the transmembrane phosphoinositide 3-phosphatase and tensin homolog 2 (*TPTE2*) gene, a paralog of the phosphatase and tensin homolog (*PTEN*) tumor suppressor [216] implicated in hepatic carcinogenesis [217] that has been found to harbor SNPs with significant allele frequency differences between males and females in European and African populations [218]. While the underlying basis for these differential autozygosity signals remains enigmatic in the absence of more detailed information on these individuals, their identification highlights the potential of our approach to identify genomic regions with differential autozygosity signals between groups that may reflect the presence of variants that have experienced differential selection histories or that influence differences in their predisposition to disease. Moreover, these findings highlight the need for further investigations among well-defined endogamic groups from India to facilitate our understanding of the genomic consequences of the long-established caste system.

**Table 4.** Genomic regions enriched for autozygosity signals in the ITU and PJL subgroups.

| Population | | | Genomic region | | | | Number of windows | Minimum $P_{perm}$ | RefSeq genes |
| --- | --- | --- | --- | --- | --- | --- | --- | --- | --- |
| ID | Name | Group | Chr | Begin (bp) | End (bp) | Length (bp) |  |  |  |
| ITU | Telugu | 1 | 10 | 132,953,074 | 133,048,305 | 95,232 | 189 | 0.013 | *TCERG1L* |
| PJJ | Punjabi | 1 | 13 | 20,001,572 | 20,181,691 | 180,120 | 28 | 0.044 | *TPTE2* |

**Figure 11.**
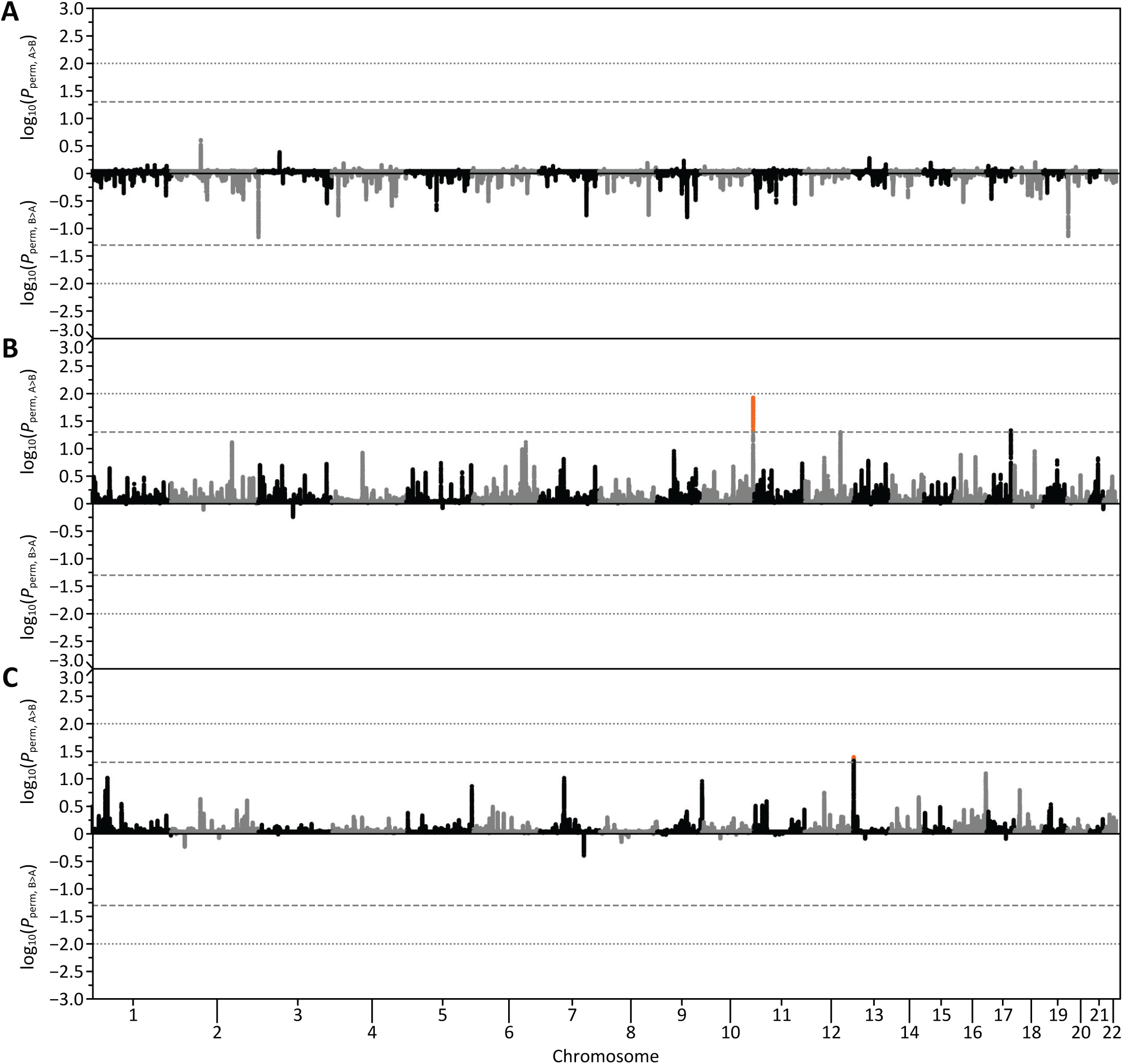
Distribution of differential ROA signals between subgroups in the GIH, ITU and PJL. Manhattan plots showing for each window the *log*_10_(*P*) of pairwise comparisons of per-individual scores in the two subgroups present in the (**A**) GIH (450 SNV window), (**B**) ITU (580 SNV window), and (**C**) PJL (610 SNV window). In each plot, *P*-values for the comparison testing whether scores in cluster A are greater than those in cluster B (Additional File 1: **Figure S22**) are shown on top with *P*-values for the reverse comparison is shown below. *P*-values represent the proportion of genome-wide maximum Wilcoxon rank-sum test statistics observed in 1,000 permutations of group labels that exceed the Wilcoxon rank-sum test statistic obtained with the true labels [150]. Windows with *P*>0.05 are shown in black and those with *P*≤0.05 are shown in orange. The horizontal grey dashed line denotes *P*=0.05 and while the grey dotted line denotes *P*=0.01.

## Discussion

We have reported an improved likelihood-based estimator for the detection of ROA in genome-wide SNV genotype data derived from either microarray platforms or WGS that accounts for autocorrelation among genotyped positions and variability in the confidence of individual genotype calls as well as the probabilities of unobserved mutation and recombination events. Fully accounting for LD among SNVs in a given window is important, because in genomic regions of high LD many pairs of individuals will share common haplotypes that are homozygous identical-by-state but not ROA in the sense defined here (i.e., inherited IBD from a common ancestor). Thus, including such spurious windows would add noise when looking for ROA for the purpose of autozygosity mapping. The incorporation of LD in our model reduces false-positive ROA detection, affording us the ability to identify smaller ROA segments with greater fidelity. An alternative approach to accounting for LD is to prune the dataset prior to its analysis. However, such an approach first requires those SNV with MAF less than 5% to be removed, which would significantly reduce the power of the *wLOD* method to detect ROA by removing those low-frequency and rare variants whose homozygosity is most indicative of autozygosity under its likelihood model (Additional File 1: **Figure S1A**). Further, such pruning cannot completely remove LD from the dataset being analyzed, with a pairwise *r*^2^ threshold of 0.5 typically applied [149]. The incorporation of LD into the model therefore better controls for the autocorrelation of autozygosity signals among nearby SNV than is attainable with LD pruning, thereby improving the specificity of the ROA it detects particularly in regions of moderate to high LD.

Similarly, accounting for the probabilities of unobserved recombination and mutation events in the genomic interval spanned by the window becomes increasingly important as a function of inter-marker distance, particularly in situations where these probabilities become nontrivial such as in lower-density microarray-derived genotype datasets. By modeling these probabilities based on the assumed number of generations since the last common ancestor of the apparent autozygous haplotypes, which we have set here based on the reported effective sizes of the populations included in The 1000 Genomes Project [155,158,161], we minimize the number of false positive ROA that can be erroneously inferred when recombination and mutations events onto very similar haplotype backgrounds give the appearance of autozygosity when paired with a non-recombined haplotype. An alternative approach would be to set an arbitrary maximum inter-marker distance allowed when calling ROA; dividing into two any inferred ROA that spans an inter-marker interval greater than that maximum. However, this has the potential to erroneously break-up long ROA, potentially impacting downstream analyses that use ROA length one of their filtering criteria. By incorporating mutation and recombination weightings into the *wLOD* model we therefore take a more informed and less-biased approach to this issue, thereby improving the detection of longer ROA particularly in datasets containing sparser sets of SNVs.

We have shown the *wLOD* ROA detection method to be well-powered to infer ROA in genetic datasets consistent with those generated by WGS and microarray-based genotyping. We recommend using this method together with a model-based ROA classification approach [18] based on genetic map lengths to distinguish ROA arising from population-level LD patterns on different evolutionary timescales (classes *G* = 1-4) from those arising from more recent cultural processes such as inbreeding (class *G* = 5). Our findings suggest that our detection approach is robust for analyses of as few as 10 individuals. However, model-based classification requires at least 25 individuals to provide a robust classification solution. Moreover, to ensure allele frequency and LD estimates used with the *wLOD* estimator are close to their true value in the population, at least 30 unrelated individuals should ideally be used in their estimation [219,220]. Intriguingly, our observation of trimodal *wLOD* score distributions for a subset of the 26 populations analyzed here, all known to practise both endogamy and consanguinity to varying degrees, suggests that this method may be able to distinguish autozygosity arising from different cultural processes that act on different time scales. Future work within well-defined endogamic and non-endogamic groups that practice consanguinity, as well as within simulated datasets exploring the breadth of possible isolation and inbreeding parameters observed in human populations, will be required to clarify this apparent property of the *wLOD* method and evaluate its potential human genetics applications.

Comparisons of the ROA inferred using the *wLOD* method on different microarray-derived and NGS datasets created from The 1000 Genomes Project Phase 3 WGS data suggest that long and to some extent intermediate length ROA are captured consistently by WGS and microarray-derived datasets. However, detection of shorter ROA does vary substantially among the different datasets as a consequence of the decreasing resolution and sensitivity attainable as the genome-wide density of genotyped positions decreases. An observation reflected in the notable lack of consistency between ROA inferred in the WES dataset and those identified in the WGS dataset. Nevertheless, population-genetic analyses of genomic ROA patterns among the 26 populations included in The 1000 Genomes Project on the basis of WGS data are consistent with our previous findings in the 64 worldwide populations included in the HGDP [221,222] and International HapMap Project [223] on the basis of ∼600,000 microarray-derived SNP genotypes [18]. These observations would therefore suggest that ROA studies using microarray-derived genotype data have similar power to detect genomic ROA patterns, and in particular those of longer ROA that are of interest to the disease genetic community due to their enrichment of deleterious variation carried in homozygous form [96,97], as those using WGS data.

We have compared the *wLOD* method against a commonly used naïve genotype counting method implemented in the software *PLINK*, as well as the recently reported HMM method of the *BCFtools* software package, under two demographic scenarios in which ROA will be of interest in population- and disease-genetic studies. In our genetic simulations the *PLINK* approach performed surprisingly well, potentially reflecting their relatively short duration which limited the opportunities for new mutations to arise on the IBD haplotypes that ultimately underlied ROA in the final generation. Indeed, only ∼4.01% and ∼14.36% of SNVs in our simulated datasets were *de novo* mutations not present in the founder individuals under scenarios 1 and 2, respectively, while just ∼2.14% and ∼2.91% of SNVs had MAF < 5%. Conversely, across the 26 populations in The 1000 Genomes Project Phase 3 WGS data on average 56% of SNVs had MAF < 5%. Nevertheless, the *wLOD* method had greater power to detect ROA versus *PLINK* across all SNV densities considered here. This difference reflects the very limited ability of the *PLINK* approach, which allows for only occasional missing or heterozygous genotypes when determining the status of a window to account for possible genotyping errors and mutations, to distinguish genomic regions that are homozygous-by-chance from those that are autozygous. In contrast, the *wLOD* method incorporates population allele frequency and LD estimates and an assumed genotyping error rate as well as accounts for the probabilities of unobserved mutations and recombination events when inferring the autozygosity status of a window, enabling more rigorous assessments of the possibility of genotyping errors and the loss of information caused by missing data. In addition, it provides a more precise measure of the probability that a given window is truly autozygous rather than simply homozygous by chance. Thus, the greater power of the *wLOD* method compared with *PLINK* reflects the greater number of false negative ROA expected under the naïve autozygosity model implemented in *PLINK*.

Comparisons of the *wLOD* method with the recently reported *RoH* function of *BCFtools* have consistently shown it to have improved power to detect ROA, and smaller ROA in particular, across all SNV densities considered here, which are representative of WGS and microarray-based genotyping platforms. However, false discovery rates of the *wLOD* method are slightly higher than those of *BCFtools/RoH*, wholly reflecting a more permissive placement of ROA boundaries marginally outside of their true locations as a consequence of the sliding window approach employed. While the underlying likelihood models of the *wLOD* and *BCFtools/RoH* approaches are similar, there are two aspects of the *wLOD* method that explain its higher power. First, by summing over all SNVs within a given window, the *wLOD* method is better able to detect the autozygosity signals of ROA comprised of older (shorter) haplotypes whose constituent SNVs individually provide only weak to modest autozygosity support than the pointwise HMM employed by *BCFtools/RoH*. Second, the *wLOD* method adjusts each SNV’s log-likelihood by the probabilities that no unobserved recombination and mutation events have occurred in the interval between it and the preceding SNV in the last *M* generations (equation 2), where *M* is set based on the expected time since the most recent common ancestor in an individual’s maternal and paternal lineages given the effective size of the population. *BCFtools/RoH* does not account for unobserved mutations in its inference model, and only allows for up to a single recombination event to have occurred within a given interval [154]. Thus, for longer ROA and those comprised of older haplotypes inherited IBD from an ancient ancestor, we would *a priori* expect *BCFtools/RoH* to have greater difficulty in making inferences as it will underestimate the number of recombination events that may have occurred as these haplotypes segregate in the general population. This may potentially underlie the noticeably erratic patterns observed with its power to detect ROA greater than 1.5 Mb in the higher SNV density simulated datasets (**Figure 6**).

Finally, the *wLOD* method distinguishes itself from *BCFtools/RoH* and *PLINK* through its ability to directly detect genomic regions enriched for autozygosity signals in one population or group compared with one or more others without requiring the inference of ROA first. We have applied this approach within the Gujarati (GIH), Punjabi (PJL), and Telugu (ITU) Asian Indian groups, comparing *wLOD* scores in two distinct clusters of individuals identified via multidimensional scaling of allele sharing dissimilarities (Additional File 1: **Figure S21**). We identified two genomic regions enriched for autozygosity signals in one of the two clusters, one in the ITU and another in the PJL, that contain genes implicated in the regulation of metabolism and the risk for developing liver cancer, respectively (**Table 4**). If we instead set a more permissive threshold of *P*_perm_ ≤ 0.1 when defining enriched regions, we identify an additional seven genomic regions marginally enriched for autozygosity in one cluster compared with the other (Additional File 2: **Table S7**). One of the seven regions was identified on chromosome 2 in ITU cluster A and contains two genes: *G6PC2*, a pancreatic glucose-6-phosphatase implicated in the modulation of fasting plasma glucose levels [224] that is a major target of cell-mediated autoimmunity in diabetes [225], and the ATP-binding cassette transporter gene *ABCB11*, mutations in which cause autosomal recessive progressive familial intrahepatic cholestasis [226,227]. In addition, a region on chromosome 17 also identified in ITU cluster A contains seven genes that include *USH1G*, mutations in which cause autosomal recessive deafness in both humans [228,229] and mice [230,231]. Finally, a region on chromosome 16 identified in PJL cluster A contains four genes including the mechanically-activated ion channel gene *PIEZO1*, mutations in which cause autosomal recessive generalized lymphatic dysplasia [232,233] as well as autosomal dominant hemolytic anemia [234,235].

The presence of genes that cause autosomal recessive diseases in three of the seven marginally significant regions—a highly unlikely observation (*P*<0.008 across 1,000 random draws of genomic regions of equivalent size)—suggests the intriguing possibility that, if these clusters do indeed represent distinct endogamic communities, they may be the hallmark of cultural and selection processes related to the differential presence of deleterious genetic variants in these genes. Future comparative autozygosity analyses of well-defined endogamic communities within the different subpopulations of India considering much larger sample sizes than were available here will facilitate our understanding of the genomic consequences of the long-established caste system and further clarify its potential role in contributing to genetic predisposition in complex disease risk and negative health outcomes.

## Conclusions

To facilitate community adoption of the *wLOD* ROA detection method as well as classification based on genetic map length via a Gaussian mixture model, we have implemented these approaches in the software *GARLIC* (G*enomic* A*utozygosity* R*egions* L*ikelihood-based* I*nference and* C*lassification*) [178] that can be downloaded at https://github.com/szpiech/garlic. As a guide, analysis of the 97 individuals in the CEU population on a Dell Precision T7600 workstation running RedHat Enterprise Linux (v.7.3) with multi-threading support enabled (16 2.60 GHz threads total) took ∼2½ minutes for the OmniExp dataset, ∼6½ minutes for the Omni2.5 dataset, and ∼40 minutes for the WGS dataset, and occupied at most ∼3 Gb, ∼7 Gb, and ∼20 Gb of RAM, respectively. Future enhancements planned for *GARLIC*’s core engine are expected to significantly reduce its runtime and memory usage. We also provide a searchable online database of ROA identified in The 1000 Genomes Project Phase 3 populations as well as a ROA genome browser based on the *JBrowse* browser interface [236] in which to explore their genomic distribution with respect to various genomic features and properties available at <link will be added during post-initial-review revision>.

## Methods

### Genotype datasets

Release v5a of Phase 3 of The 1000 Genomes Project (accessed March 29^th^, 2015) provides phased genotypes at 84,801,880 genetic variants in 2,504 individuals from 26 worldwide human populations discovered using a low-coverage WGS approach [155]. During the genotype phasing, occasional positions with missing genotypes were imputed; consequently, our datasets contain no missing data. We first developed a subset of this WGS dataset in which to perform individual-level quality control prior to developing different subsets in which to evaluate the performance of the *wLOD* method. In all subsets we applied a common set of quality-control procedures described in Pemberton *et al*. [237] to remove low-quality variants (Additional File 1: **Figure S22**).

#### Individual-level quality control

To independently verify the putative unrelatedness and population labeling of individuals reported by The 1000 Genomes Project Consortium, we developed a preliminary Omni dataset comprised of the 2,165,831 autosomal, 48,458 X-chromosomal, and 543 Y-chromosomal SNPs in The 1000 Genomes Project data that are present on the Illumina HumanOmni2.5-8 BeadChip (stage 1; Additional File 1: **Figure S22**). Across the 1,693 individuals for which genotypes derived using the HumanOmni2.5-8 BeadChip were also available, genotype concordance between the WGS- and BeadChip-derived genotypes lay between 0.99431 and 0.99986 (mean=0.99953, SD=0.00041). We identified intra- and inter-population pairs of individuals related closer than first cousins as well as those individuals whose reported sex or population labels were likely to be erroneous as described in Pemberton *et al*. [237]. Using these approaches, we identified six individuals whose reported sex is likely to be erroneous, 47 individuals who did not cluster genetically with other individuals sharing the same population label, and 14 intra-population and one inter-population pairs of close relatives (Additional File 2: **Table S8**).

#### Preparation of final datasets

Removing one individual from each intra-population relative pair, both individuals from the inter-population relative pair, and the 53 individuals whose reported sex or population labels were suspected to be erroneous (68 total individuals; Additional File 2: **Table S8**), we developed four subsets of The 1000 Genomes Project data that were restricted to the 2,436 unrelated individuals and autosomal biallelic variants (stage 2; Additional File 1: **Figure S22**).

First, we developed a WGS dataset comprised of 75,071,695 SNVs. Second, we developed a WES dataset comprised of the 1,830,512 SNVs that are present within the regions captured by the Roche Nimblegen SeqCap EZ Human Exome Library v3.0 system. Third, we developed an Omni2.5 dataset comprised of the 2,166,414 SNPs that are present on the Illumina HumanOmni2.5-8 BeadChip. Fourth, as ∼96% of all markers present on the Illumina HumanOmniExpress-24 BeadChip are also present on the HumanOmni2.5-8 BeadChip, we developed an OmniExpress dataset comprised of the 676,445 SNPs in the Omni2.5 dataset that are present on the HumanOmniExpress-24 BeadChip.

#### Geographic distances

The geographic distance of each population from Addis Ababa, Ethiopia, was calculated as in Rosenberg *et al*. [238] with the use of waypoint routes, based on the sampling location reported by The 1000 Genomes Project [155].

### Simulation of genetic datasets

For two demographic scenarios, we generated 50 independent replicates of genetic datasets using a forward-in-time process as previously described [176]. In their original approach, prior to performing the simulation steps Kardos *et al*. placed *N* predetermined polymorphic SNV onto the chromosome’s genetic map by randomly sampling *N* unique genetic map positions in the range 0 to *g*_*max*_ (the user-defined genetic map length of the simulated genome), only converting genetic map positions to physical map positions based upon a fixed user-defined recombination rate to physical map distance relationship when writing the simulated datasets to file. Here, we modified their approach to instead create a non-uniform distribution of recombination rates across the simulated chromosome and allow any base pair to mutate during the simulation.

If we let *g*_*p*_ represent the genetic map position assigned to physical map position *p*, which is equal to the base pair count from the beginning of the chromosome. Based on the user-defined values for *g*_*max*_ and recombination rate θ, all values of *g* lie within the interval [0, *g*_*max*_] and all values of *p* lie within the interval [1 .. (*g*_*max*_/θ)×1,000,000]. To begin, we created a backbone of genetic and physical map positions onto which we will place all other positions, randomly drawing (*g*_*max*_/θ) + 1 values of *g* and assigning them in increasing order to *p* in the range [1 .. (*g*_*max*_/θ)] (i.e. every Mb). Next, we randomly chose *N* values of *p* to be predetermined polymorphic SNVs, and then randomly assigned each a value of *g* based upon the backbone interval in which it was located, again ensuring that values of *g* always increase as a function of *p*. Finally, all values of *p* that were not among the set of predetermined SNVs were assigned a value of *g* through interpolation onto the construct created by the values of *p* and *g* assigned to the predetermined SNVs. This approach created a non-uniform relationship between physical and genetic map distance along the simulated chromosome that is similar to that observed on real human chromosomes (not shown).

To extend the method of Kardos *et al*. to enable any base pair on the simulated chromosome to mutate, for each individual in each generation, the number of mutations that occur during each meiosis was drawn from a Poisson distribution with mean μ×[(*g*_*max*_/θ)×1,000,000], where *μ* is mutation rate. The base pairs to be mutated were then chosen at random from all (*g*_*max*_/θ)×1,000,000 possible positions without replacement. Mutations were tracked and then incorporated into the genotypes of individuals in the analyzed dataset; all monomorphic positions were removed during dataset construction.

In all simulations, we set *g*_*max*_ to 325 cM, θ to 1.3 cM/Mb [239], and μ to 1.18×10^−8^ [160], and scaled θ and μ by a factor of 10 to increase genetic diversity in the final generation [240]. *N* was chosen separately for each simulated scenario such that the final number of polymorphic SNVs in the dataset (both predetermined and *de novo*) was ∼750,000; *N*=725,000 for scenario 1 and *N*=650,000 for scenario 2. Because predetermined polymorphic SNVs can become fixed over the course of the simulation, their numbers in the analyzed datasets lay between 679,256-717,855 (25,788-31,503 *de novo* SNVs) for scenario 1 and between 633,582-638,675 (103,871-110,077 *de novo* SNVs) for scenario 2.

### Calculation of *LOD and wLOD* estimators

To minimize the number of variables that varied in within-dataset comparisons, we used a single set of allele frequencies when calculating *wLOD* and *LOD* scores at all window sizes considered. To account for sample-size differences among populations, we used a resampling procedure to estimate the allele frequencies, sampling 100 non-missing alleles with replacement and calculating allele frequencies from these 100 alleles. As a consequence of the resampling procedure, it was possible for an individual to possess an allelic type whose frequency was estimated to be 0 in the sample of 100 alleles. SNV positions at which this scenario was encountered were treated as missing when calculating *wLOD* and *LOD* scores for all windows containing the positions in individuals that had the allelic type of frequency 0.

As our datasets contained phased genotypes, in the LD correction applied in the *wLOD* estimator (equation 3) LD was estimated with the correlation coefficient *r*^2^ [241] using a resampling procedure to account for the possible influence of sample size on homozygosity-based LD statistics [219]. For each pair of SNPs, we randomly sampled 55 individuals-the smallest population sample size in our dataset (**Table 2**)–without replacement and the LD computation was performed using those 55 individuals. Note that we used a single set of LD estimates when calculating *wLOD* scores at all window sizes considered.

In the recombination rate correction applied in the *wLOD* estimator (equation 4), the genetic map position of each marker in the Omni2.5 dataset and its subsets were downloaded from the Laboratory of Computational Genetics at Rutgers University (http://compgen.rutgers.edu). The genetic map position of each marker in the WES and WGS datasets was determined by interpolation onto the Rutgers linkage-physical map [242] based on their UCSC Build hg19 physical map position.

Due to computer memory requirements for Gaussian kernel density estimation, the *wLOD* score distributions used to determine the autozygosity score thresholds in the WGS dataset considered only twenty individuals chosen at random. Based on our investigation into the effect of sample size on score threshold (Additional File 1: **Figure S10**), we do not expect this approach to have biased our detection of ROA in the WGS dataset. All genome-wide windows were, however, considered when determining optimal window sizes in the Omni2.5, OmniExpress, and WES datasets.

### Classification of ROA

We ran unsupervised Gaussian fitting of the ROA length distribution using the *mclust* package (v.5.2) [243] in *R* v.3.3.3 [244], allowing component magnitudes, means, and variances to be free parameters. BIC likelihoods with increasing number of components (*G*) were calculated using the function *mclustBIC*, while final classification under the five component model was performed using the function *Mclust*.

### Genomic distribution and geographic patterns of ROA

The frequency at which each SNV was present in ROA in each population was calculated as described in Pemberton *et al*. [18]. To compare the genomic distribution of ROA across populations, we calculated mean ROA frequencies in non-overlapping 50 kb windows across all SNVs polymorphic in that population that were within the window, and excluding windows that lay within the centromere and telomeres. To evaluate the similarity of ROA frequency patterns among populations, we performed classical (metric) multidimensional scaling (MDS) separately for each ROA length class based on a matrix of ROA frequency dissimilarities between all pairs of populations, calculated as one minus the Pearson correlation coefficient (*r*) of their mean ROA frequencies across windows. We then applied MDS to this matrix using *cmdscale* in *R*.

We compared population patterns in the MDS based on ROA frequencies to an MDS based on a matrix of pairwise *F*_ST_ among populations calculated with our WGS dataset and the method of Hudson *et al*. [245] according to the recommendations of Bhatia *et al*. [246]. The similarity of patterns in our MDS of ROA dissimilarities and those in the MDS of *F*_ST_ was evaluated with the Procrustes method [247].

### Relationship between ROA and genomic variables

For each ROA length class, we investigated recombination rate and haplotype-based *nS*L selection scores [206] for correlations with ROA frequency across the autosomes. Population-based recombination-rate estimates were obtained from Phase 3 of The 1000 Genomes Project [155] (downloaded July14^th^, 2014), and *nS*L values for each of the 26 populations were calculated in the WGS dataset considering only SNVs with MAF > 0.05 and normalization of unstandardized scores in 100 genome-wide frequency bins with *selscan* [248]. Comparisons between ROA frequency and recombination rate and *nS*_L_ were performed as described in Pemberton *et al*. [18] considering the mean value of each variable in non-overlapping 50 kb windows, excluding windows within the centromere and telomeres, calculated across all SNV within the window for which the variable was available. Admixed Afro-European (ASW and ACB) and Mestizo (CLM, MXL, PEL, and PUR) populations and the geographically imprecise CEU (Utah residents of Northwestern European ancestry) group were omitted from geographic analyses but were included in the scatterplots.

### List of Abbreviations Used

ASW, African American; ACB, Afro-Caribbean; BEB, Bengali; bp, base-pair; CEU, European American; CDX, Dai; CHB, Northern Han; CHS, Southern Han; CLM, Colombian; cM, centimorgan; ESN, Esan; FIN, Finnish; GIH, Gujarati; GWD, Gambian; GBR, British; HGDP, Human Genome Diversity Panel; HMM, hidden Markov model; IBD, identical by descent; IBS, Iberian; Indel, insertion/deletion variant; ITU, Telugu; JPT, Japanese; kb, kilobase; KHV, Kinh; LD, linkage disequilibrium; *LOD*, logarithm of the odds; *wLOD*, weighted logarithm of the odds; LWK, Luhya; MAF, minor allele frequency; Mb, megabase; MSL, Mende; MXL, Mexican American; NGS, next-generation sequencing; OMIM, Online Mendelian Inheritance of Man; PEL, Peruvian; PJL, Punjabi; PUR, Puerto Rican; RAM, random access memory; ROA, regions of autozygosity; ROH, runs of homozygosity; SNP, single-nucleotide polymorphism; SNV, single-nucleotide variant; STU, Sri Lankan Tamil; TSI, Toscani; WES, whole-exome sequencing; WGS, whole-genome sequencing; YRI, Yoruban.

## Declarations

### Ethics approval and consent to participate

This study was approved by the University of Manitoba Health Research Ethics Board (protocol ID H2013:141).

### Consent for publication

All authors read and approved the final manuscript.

### Availability of data and material

The raw genetic data analyzed during the current study are available in The International Genome Sample Resource repository, http://www.internationalgenome.org/data. The software *GARLIC* implementing the new method reported by this study can be obtained at https://github.com/szpiech/garlic (v1.1.0 and later).

## Authors’ contributions

T.J.P. conceived the study. T.J.P and Z.A.S. developed the weighted likelihood estimator. T.J.P prepared the datasets and performed the genetic simulations with the assistance of Z.A.S. A.B. and M.K. performed the comparative analyses. A.B. performed the genomic analyses with the assistance of T.J.P. Z.A.S. implemented the *wLOD* method in the software *GARLIC*. A.B. and T.J.P. wrote the paper with the assistance of Z.A.S. and M.K.

## Acknowledgements

The authors would like to thank Noah Rosenberg for useful comments and discussions.

## Description of Additional Data Files

**Additional File 1:** Figures S1-22

**Additional File 2:** Tables S1-8

